# Neutral Frustration Landscape Architecture of SARS-CoV-2 Spike-Antibody Interfaces Shapes Immune Evasion Mechanisms for Ultrapotent Neutralizing Antibodies and Determines Pathways of Viral Adaptation: Insights from Integrative Computational Approach

**DOI:** 10.1101/2025.10.31.685956

**Authors:** Mohammed Alshahrani, Vedant Parikh, Brandon Foley, Gennady Verkhivker

## Abstract

The relentless evolution of SARS-CoV-2 underscores the urgent need to decipher the molecular principles that enable certain antibodies to maintain exceptional breadth and resilience against immune escape. In this study, we employ a multi-pronged computational framework integrating structural analysis, conformational dynamics, mutational scanning, MM-GBSA binding energetics, and conformational and mutational frustration profiling to dissect the mechanisms of ultrapotent neutralization by a cohort of broadly reactive Class 1 antibodies (BD55-1205, 19-77, ZCP4C9, ZCP3B4) and the Class 4/1 antibody ADG20. We reveal a unifying biophysical architecture: these antibodies bind via rigid, pre-configured interfaces that distribute binding energy across extensive epitopes through numerous suboptimal yet synergistic interactions, predominantly with backbone atoms and conserved side chains. This distributed redundancy enables tolerance to mutations at key sites like F456L or A475V without catastrophic loss of affinity. Mutational scanning identifies a hierarchical hotspot organization where primary hotspots (e.g., H505, Y501, Y489, Y421) which overlap with ACE2-contact residues and incur high fitness costs upon mutation are buffered by secondary hotspots (e.g., F456, L455) that are more permissive to variation. MM-GBSA energy decomposition confirms that van der Waals-driven hydrophobic packing dominates binding, with primary hotspots contributing disproportionately to affinity, while electrostatic networks provide auxiliary stabilization that mitigates mutational effects. Critically, both conformational and mutational frustration analyses demonstrate that immune escape hotspots reside in neutral-frustration landscape that permit mutational exploration without destabilizing the RBD, explaining the repeated emergence of convergent mutations across lineages. Our results establish that broad neutralization arises not from ultra-high-affinity anchors, but rather from strategic energy distribution across rigid, evolutionarily informed interfaces. By linking distributed binding, neutral frustration landscapes, and viral fitness constraints, this framework provides a predictive blueprint for designing next-generation therapeutics and vaccines capable of withstanding viral evolution.

## Introduction

The emergence of severe acute respiratory syndrome coronavirus 2 (SARS-CoV-2) has catalyzed an extraordinary global research response focused on unraveling its molecular structure, mechanisms of host cell entry, and the immune responses it provokes. At the heart of these studies is the SARS-CoV-2 Spike (S) glycoprotein—a trimeric surface protein essential for viral entry and the principal target of neutralizing antibodies.^1–15^ The S protein displays remarkable conformational flexibility, allowing it to transition through multiple functional states—from initial receptor binding to membrane fusion—while evading host immune detection.^1–15^ Structurally, it consists of two functionally distinct subunits: S1 and S2. The S1 subunit comprises four key domains: the N-terminal domain (NTD), the receptor-binding domain (RBD), and two conserved subdomains, SD1 and SD2—each playing a specialized role in the virus’s life cycle.^16–18^ The SD1 and SD2 subdomains play essential structural roles in maintaining the prefusion conformation of the S protein, acting as molecular scaffolds that regulate the timing and efficiency of membrane fusion.^16–18^ The structural variability introduced by mutational changes of SARS-CoV-2 S protein variants of concern (VOCs) can modulate the virus capacity to bind to the ACE2 receptor and evade immune responses, complicating the host’s ability to mount an effective defense. Extensive cryo-electron microscopy (cryo-EM) and X-ray structures of SARS-CoV-2 S protein variants together with their complexes with antibodies underscore the balance between structural stability, immune evasion, and receptor binding that shapes the evolutionary trajectory of SARS-CoV-2 and its variants.^19–25^

The continued evolution of SARS-CoV-2 within the Omicron lineage and its descendants such as XBB.1, XBB.1.5, and more recent subvariants (e.g., JN.1, KP.2, KP.3), highlights not only the virus extraordinary adaptability but also the emergence of convergent evolutionary hotspots— specific residues that are repeatedly and independently mutated across geographically and temporally distinct lineages.^26–29^ The recurrence of identical or functionally similar mutations (such as F456L, R346T, L455F, and K444T) across unrelated variants signals a narrowing evolutionary landscape, where SARS-CoV-2 is increasingly optimizing a limited set of high-impact changes to balance transmissibility, immune evasion, and structural stability.^26–29^

The JN.1-derived subvariants KP.2 and KP.3 have independently acquired a constellation of key S mutations—including R346T, F456L, Q493E, and V1104L—that collectively enhance both transmissibility and the ability to evade neutralizing antibodies. Additional offshoots of JN.1, such as LB.1 and KP.2.3, further illustrate this dynamic evolutionary pattern: they share hallmark mutations R346T and F456L, while also introducing distinct changes S:S31-(a deletion at position 31) and S:Q183H in LB.1, or S:H146Q in KP.2.3.^26–29^ Notably, the F456L substitution has emerged as a pivotal driver of immune escape, with KP.3 recognized as one of the most antibody-evasive sublineages within the JN.1 clade.^29^ This rapid diversification underscores extraordinary capacity of the virus to adapt under immune pressure, fine-tuning mutations to balance evasion with essential functional requirements such as receptor binding. Strikingly, the repeated, independent emergence of mutations at positions L455F, F456L, and R346T across unrelated lineages signals strong convergent evolution, reflecting intense selective pressure to optimize viral fitness. Supporting this, recent cryo-EM structural analyses of RBD complexes from JN.1, KP.2, and KP.3 reveal that the F456L mutation synergizes with Q493E to strengthen ACE2 binding providing a structural basis for KP.3 heightened infectivity and immune resistance.^30,31^

The recombinant variant XEC, which originated from KP.3, has recently drawn attention due to two additional mutations in the NTD F59S and T22N.^32,33^ XEC displays markedly higher infectivity than its parental KP.3 lineage and exhibits enhanced resistance to neutralizing immune responses. Separately, the KP.3.1.1 subvariant emerged from KP.3 through a deletion at position S31 in the NTD. Structural and binding analyses of the KP.3.1.1 RBD in complex with ACE2 revealed a critical epistatic interaction between two key RBD mutations F456L and Q493E.^33^ In recent months, the JN.1-derived LP.8.1 and LP.8.1.1 sublineages have now largely displaced XEC across Europe and North America. Concurrently, LF.7 and its descendant LF.7.2.1 gained prominence in Asia, while MC.10.1 experienced a sharp, albeit brief, surge in global prevalence.^34,35^ LF.7 carries seven additional Spike mutations relative to JN.1: four in the NTD: T22N, S31P, K182R, and R190S—and three in the RBD – R346T, K444R, and F456L [34,35]. Its successor, LF.7.2.1 variant acquired an additional A475V substitution in the RBD and quickly outcompeted LF.7 in Asian regions (Figure 1, Supporting Information, Table S1).

**Figure 1.**
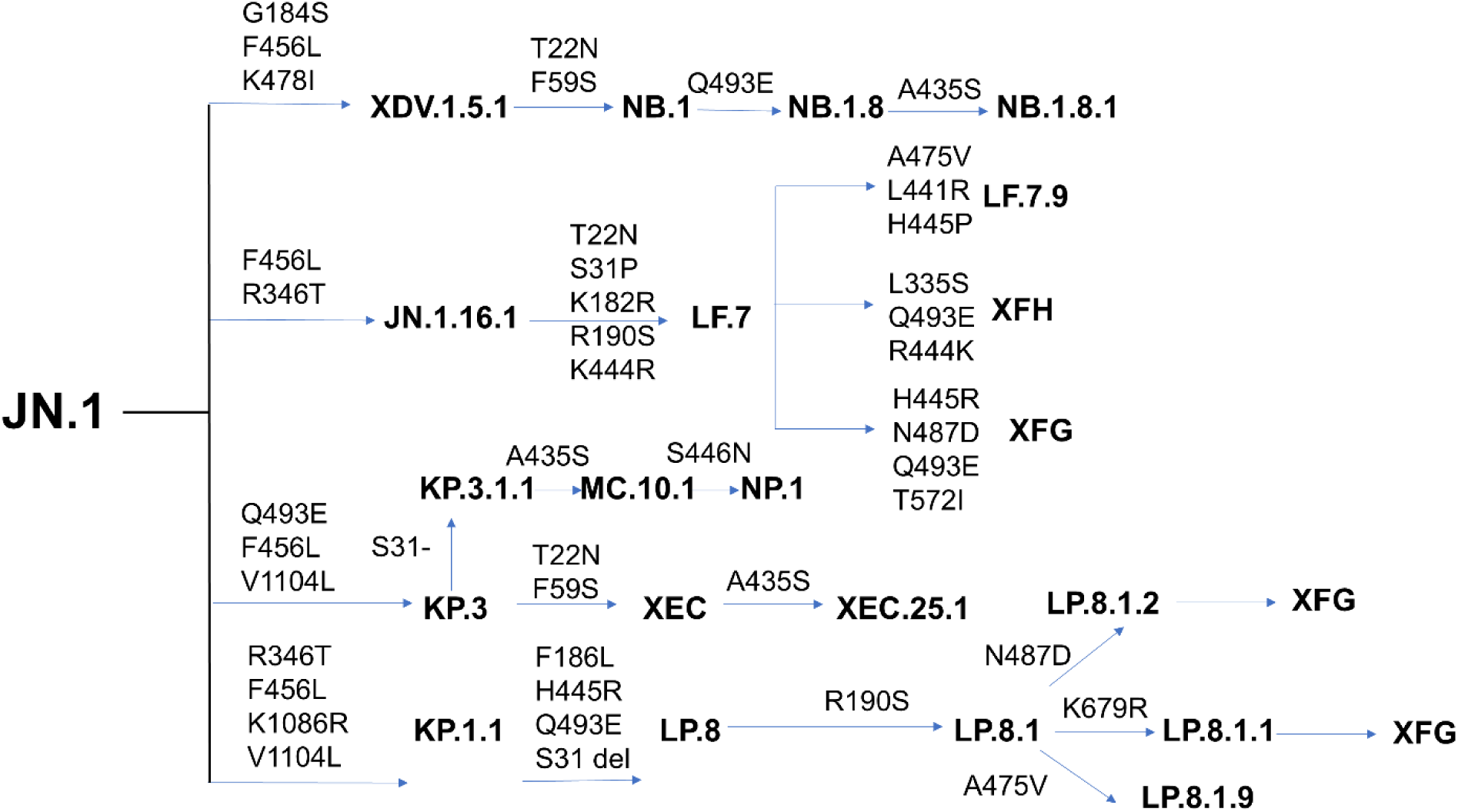
The evolutionary trajectories of the SARS-CoV-2 variants originated from JN.1 with the main branches of Omicron variants and their respective mutations.

By June 2025, two newer variants NB.1.8.1 and the recombinant XFG began showing a pronounced growth advantage over earlier JN.1 sublineages on a worldwide scale. NB.1.8.1, emerging from the recombinant lineage XDV, harbors seven S mutations beyond those in JN.1, namely T22N, F59S, and G184S in the NTD, and A435S, F456L, K478I, and Q493E in the RBD.^36,37^ XFG—a recombinant offspring of LF.7 and LP.8.1.2 currently under active monitoring—carries four additional mutations beyond LF.7: H445R, N487D, and Q493E in the RBD, and T572I in the SD1.^36,37^ Together, these variants illustrate the accelerating pace and increasing complexity of SARS-CoV-2 evolution, driven by convergent mutations and recombination events that enhance transmissibility and immune escape (Figure 1)

A cornerstone of the immune response to SARS-CoV-2 is the generation of antibodies directed against the viral S protein and particularly the RBD which is essential for host cell entry.^38,39^ Advances in high-throughput yeast display screening and deep mutational scanning (DMS) have enabled precise mapping of antibody escape mutations across the RBD and the systematic classification of neutralizing antibodies into six major epitope groups (A–F) based on their binding footprints.^38,39^ To dissect the molecular basis of broadly neutralizing antibodies elicited by XBB/JN.1 infections, Cao and colleagues employed large-scale yeast-display DMS assays, profiling escape mutations across a panel of 2,688 monoclonal antibodies, including 1,874 isolated from individuals convalescing from XBB/JN.1 infection.^40^ By integrating these high-throughput DMS datasets, they engineered pseudoviruses bearing predicted escape mutations and used them as selective filters to identify antibodies resilient to viral evolution, demonstrating that future RBD evolutionary trajectories and immune escape trends could be programmed. In a most recent groundbreaking study, Cao’s team harnessed DMS technology not only to anticipate SARS-CoV-2 evolutionary trajectories but also to strategically isolate monoclonal antibodies capable of neutralizing both current and future viral variants. A retrospective analysis of 1,103 monoclonal antibodies originally derived from wildtype (WT) SARS-CoV-2 infection demonstrated the power of this approach as it boosted the likelihood of identifying antibodies effective against the XBB.1.5 variant from a mere 1% to 40%.^41^ Using this predictive screening framework, this study discovered class 1 antibody BD55-1205 which is the only WT-elicited monoclonal antibody to date with pan-variant neutralizing activity. BD55-1205 displays remarkable breadth, potently neutralizing all major SARS-CoV-2 lineages, including XBB, BA.2.86, and diverse JN.1-derived subvariants, as well as computationally predicted escape mutants.^41^ DMS studies showed that BD55-1205 may be affected by mutations at several RBD sites in the range from 503-505, and also by mutations at sites F456, A475, and Q493, all of which interact with the host cell receptor ACE2.^42^ While mutations changes in F456 and A475 sites are deleterious for ACE2 binding, A475V mutation escapes BD55-1205 and is only mildly deleterious for ACE2 binding, recently emerging in several JN.1-descendant lineages (Figure 1).^41,42^ Another class 1 broadly neutralizing antibody VIR-7229 demonstrated potent neutralizing activity against a broad spectrum of circulating variants, including EG.5, BA.2.86, and JN.1.^43^ Structural analyses of VIR-7229 in complex with the XBB.1.5 and EG.5 RBD revealed its remarkable adaptability as the antibody effectively engages the RBM with F456 or L456 at the key site of immune-driven variation. This flexibility allows VIR-7229 to tolerate extensive epitope diversity while maintaining high-affinity binding and inducing high barrier to developing resistance.^43^ Our preliminary computational analysis suggested the unique binding resilience of the BD55-1205 antibody to immune escape may arise by leveraging a broad epitope footprint and distributed hotspot architecture, additionally supported by backbone-mediated specific interactions, which together enable exceptional tolerance to mutational escape.^44^ A computational study of VIR-7229 antibody binding to XBB.1.5 and EG.5 Omicron variants also emphasized a large and structurally complex epitope, demonstrating certain structural adaptability and compensatory energetic effects to F456L and L455S mutations.^45^ Our preliminary computational studies suggested that these class I antibodies could synergistically exploit a combination of different factors including broad epitope coverage, reliance on multiple binding hotspot residues, and energetic tolerance to mutations with subtle interaction differences that could determine the unique neutralization profiles and resilience to immune escape across a wide range of new Omicron variants.^44,45^

Class 1 human monoclonal antibody 19-77 demonstrates potent neutralizing activity against the majority of SARS-CoV-2 variants.^46^ The engineeother broadlyned variants of 19-77 antibody – 19-77R71A, 19-77R71L, and 19-77R71V – achieved neutralization across 32 variants tested, including the currently dominant KP.3.1.1 and XEC lineages.^46^ The improved activity of optimized 19-77 antibodies was attributed to the loss of the internal hydrogen bonding by R71 leading to greater flexibility of the CDRs which in turn, allowed the optimized antibodies to be more tolerant of mutations within the epitope, ultimately resulting in the broad neutralization potency and breadth.^46^ It was noted however, that the applicability of this optimization strategy did not extend to other SARS-CoV-2-neutralizing antibodies suggesting a specific mutation is not likely to confer a generalizable benefit to other broadly neutralizing monoclonal antibodies. ZCP4C9 and ZCP3B4 are other recently discovered class 1 broadly neutralizing human monoclonal antibodies that target the RBD epitopes extensively overlapping with the binding site of ACE2 and showing exceptional ability to neutralize a wide range of SARS-CoV-2 variants—including early strains and highly immune-evasive Omicron sublineages.^47^ The structural footprints of ZCP3B4 and ZCP4C9 avoided most of the mutation sites in RBD and did not contain any convergent mutations carried by BQ.1.1, XBB, EG.5.1 or BA.2.86 sublineages to minimize the impact of immune escape.^47^ In general, this growing but still exclusive cohort of ultrapotent and broadly neutralizing class 1 human monoclonal antibodies revealed the power of directly competing with ACE2 binding site and exploiting the diversity of the RBD interaction mechanisms to successfully combat the full spectrum of circulating variants.

While most of clinically approved monoclonal antibodies have been rendered largely ineffective by the antigenic drift of Omicron sublineages to the evolving JN.1 subvariants, ‘class 4 (group F3) antibodies pemivibart (VYD222)^48^ and SA55 (BD55-5514)^49^ showed remarkable neutralization breath against newest JN.1 descendant variants and target a similar RBD epitope and demonstrate broad neutralization against SARS-CoV-2 variants. SA55 effectively prevents viral escape by JN.1 sublineages, likely due to its substantially increased binding affinity to the virus where the efficacy of SA55 is affected by Y508H, moderately affected by G504S, strongly affected by K440E, and escaped only by V503E and G504D mutations.^49^ VYD222 is a human re-engineered antibody derived from ADG20 which extends its binding footprint from the RBD site to the highly conserved site on the RBD known as CR3022 site.^50^ The crystal structures of VYD222 complexed with the RBD of prototype SARS-CoV-2 and Omicron BA.5 demonstrated the binding epitope spans from the receptor binding site to the conserved CR3022 site, while DMS results indicated that mutations of epitope residues 403-405, 407-409, 436-437, and 500-506 can induce antibody escape, but most of these mutations affect the host receptor binding, which may therefore decrease viral fitness.^51^ A comparative analysis of SA55 and VYD222 escape mutations revealed shared features including mutations in the NTD, within the conserved region on the epitope and a shared V503E mutation, generally revealing that escape mutations are rare in currently circulating SARS-CoV-2 populations.^52^

The interplay of dynamic and energetic factors is central to understanding how immune escape hotspots evolve, as these adaptations allow the virus to evade neutralizing antibodies without compromising its infectivity. Computer simulations have significantly advanced our understanding of the dynamics and functions of the S complexes with ACE2 and antibodies at the atomic level.^53–55^ Conformational dynamics and allosteric interactions can be linked to binding of novel human antibodies^56,57^ where antibody-induced associated changes in S dynamics can distinguish weak, moderate and strong neutralizing antibodies.^58^ Recent computational and structural studies suggested a mechanism in which the pattern of specific escape mutants for ultrapotent antibodies may be driven by a complex balance between the impact of mutations on structural stability, binding strength, and long-range communications.^59,60^

In the present study, we considerably expanded the scope, the computational framework and the systematic analysis of the binding, evolution and immune escape mechanisms for ultrapotent Class I neutralizing antibodies by adding to previously examined BD55-1205 and VIR-7229 antibodies^44,45^, some very recently unveiled 19-77 antibody in complexes with RBD nd HK.3 variants, 19-77 R71 mutants, ZCP3B4 and ZCP4C9 antibody in complexes with BA.5 RBD. For comparison, we provide a head-to-head analysis for another broadly neutralizing class 4/1 ADG20 antibody that targets a much smaller epitope of conserved RBD residues. We employ a multi-pronged computational modeling approach which combined coarse-grained simulations and atomistic reconstruction of conformational landscapes together with mutational scanning of the binding interfaces, prediction of immune escape and binding hotspots, rigorous binding free energy analysis and the energy landscape-based global frustration analysis of adaptive evolution. Here, we systematically analyze and critically examine two major binding scenarios that were put forward as potential mechanisms underlying activity of these class 1 broadly neutralizing antibodies: conservation-driven binding, where antibodies exploit highly conserved residues critical for viral function, and adaptability-driven binding where antibodies utilize structural flexibility and compensatory interactions to tolerate mutations while maintaining neutralization efficacy. We show that these antibodies employ distinct binding mechanisms, targeting unique epitopes on the RBD with varying degrees of conservation and flexibility. Conformational dynamics and mutational scanning analyses reveal that different antibodies employ diverse molecular strategies to engage the RBD, shaped by their distinct structural architectures and chemical complementarity. To examine how these balancing relationships between structural stability and plasticity we quantified conformational and mutational frustration^61–64^ of S protein residues using the equilibrium conformational distributions. Our results show that the dominant pattern of neutral frustration in the evolutionary hotspots on the RBD may create a state of “energetically suboptimal, adaptable frustration” that limits access to potentially superior adaptive outcomes but generates mutational “hotspots” – genomic regions where mutations repeatedly and predictably occur, leading to highly repeatable evolutionary trajectories. The results of this study demonstrate that broad neutralization can emerge not from a few ultra-high affinity “anchor” residues, but from a rigid, pre-configured interface that distributes binding energy across a broad epitope surface via numerous suboptimal, yet synergistic, interactions. A comparison with class 4/1 ADG20 antibody illustrates a considerable generality of conservation-driven binding mechanism where recognition of a minimally frustrated epitope can induce allosteric hypermobility in the distal RBM loop, a region already prone to immune-driven mutations.

The discovered relationships indicate that neutral frustration at the key adaptable evolutionary hotspots of the RBD-antibody interfaces can create mutational path leading to rapid adaptation of immune escape mutations that represent suboptimal and yet robust recurrent outcomes. This framework also explains the emergence of recent variants: mutations such as F456L, A475V, and L455F are selected from a limited set of viable escape routes permitted by the neutral-frustration architecture of the RBM. Overall, this study demonstrates that the evolutionary trade-offs SARS-CoV-2 faces—balancing immune evasion against the need to maintain ACE2 binding—are often based on energetically neutral, suboptimal variations that are highly context-dependent even for the same class of broadly neutralizing antibodies The emerging understanding of the immune evasion mechanisms suggests that at the molecular level, the evolution of immune escape hotspots in SARS-CoV-2 which is a complex process can be influenced by both random drift of mutations due to neutrally frustrated architecture and natural selection that acts on these variants, favoring convergent evolution for those positions that provide a recurrent fitness advantage.

## Materials and Methods

### Structure Preparation and Analysis

All structures were obtained from the Protein Data Bank.^65^ Hydrogen atoms and missing residues were initially added and assigned according to the WHATIF program web interface.^66,67^ The structures were further pre-processed through the Protein Preparation Wizard (Schrödinger, LLC, New York, NY) for assignment and adjustment of ionization states, formation of assignment of partial charges as well as additional check for possible missing atoms and side chains that were not assigned by the WHATIF program. The missing loops in the cryo-EM structures were reconstructed using template-based loop prediction approaches ModLoop^68^ and ArchPRED^69^ and further confirmed by FALC (Fragment Assembly and Loop Closure) program.^70^ The side chain rotamers were refined and optimized by SCWRL4 tool.^71^ The protein structures were then optimized using atomic-level energy minimization using 3Drefine method.^72^

### Coarse-Grained Brownian Dynamics Simulations

Although all-atom MD simulations with the explicit inclusion of the glycosylation shield can in principle provide a rigorous assessment of conformational landscape of the SARS-CoV-2 S proteins, such direct simulations may become challenging due to the size of a complete SARS-CoV-2 S system embedded onto the membrane and this complexity can often obscure the main molecular determinants of the binding mechanisms and prevents efficient comparative analysis across large number of antibodies. Coarse-grained (CG) models are computationally effective approaches for rapid and efficient exploration of large systems over long timescales.

Coarse-grained Brownian dynamics (BD) simulations have been conducted using the ProPHet (Probing Protein Heterogeneity) approach and program.^73–75^ BD simulations are based on a high resolution CG protein representation^76^ of the SARS-CoV-2 S Omicron trimer structures that can distinguish different residues. In this model, each amino acid is represented by one pseudo-atom at the Cα position, and two pseudo-atoms for large residues. The interactions between the pseudo-atoms are treated according to the standard elastic network model (ENM) in which the pseudo-atoms within the cut-off parameter, *R*_c_ = 9 Å are joined by Gaussian springs with the identical spring constants of γ = 0.42 N m^−1^ (0.6 kcal mol^−1^ Å^−2^. The simulations use an implicit solvent representation via the diffusion and random displacement terms and hydrodynamic interactions through the diffusion tensor using the Ermak-McCammon equation of motions and hydrodynamic interactions as described in the original pioneering studies that introduced Brownian dynamics for simulations of proteins.^77,78^ We adopted Δ*t* = 5 fs as a time step for simulations and performed 100 independent BD simulations for each system using 100,000 BD steps at a temperature of 300 K.

### Coarse-Grained CABS Simulations and Atomistic Reconstruction of Ensembles

CABS-flex approach is employed that efficiently combines a high-resolution coarse-grained model and efficient search protocol capable of accurately reproducing all-atom MD simulation trajectories and dynamic profiles of large biomolecules on a long time scale.^79–84^ In this high-resolution model, the amino acid residues are represented by Cα, Cβ, the center of mass of side chains and another pseudoatom placed in the center of the Cα-Cα pseudo-bond. In this model, the amino acid residues are represented by Cα, Cβ, the center of mass of side chains and the center of the Cα-Cα pseudo-bond. The CABS-flex approach implemented as a Python 2.7 object-oriented standalone package was used in this study to allow for robust conformational sampling proven to accurately recapitulate all-atom MD simulation trajectories of proteins on a long time scale. Conformational sampling in the CABS-flex approach is conducted with the aid of Monte Carlo replica-exchange dynamics and involves local moves of individual amino acids in the protein structure and global moves of small fragments. The default settings were used in which soft native-like restraints are imposed only on pairs of residues fulfilling the following conditions: the distance between their *C*_α_ atoms was smaller than 8 Å, and both residues belong to the same secondary structure elements. A total of 1000 independent CG-CABS simulations were performed for each of the systems studied. In each simulation, the total number of cycles was set to 10,000 and the number of cycles between trajectory frames was 100. MODELLER-based reconstruction of simulation trajectories to all-atom representation^85^ provided by the CABS-flex package was employed to produce atomistic models of the equilibrium ensembles for studied systems. The root mean square deviations (RMSD) between the reference state and a trajectory of conformations were computed using MDTraj Python library^86,87^ and MDAnalysis Python toolkit (www.mdanalysis.org) utilizing the fast QCP algorithm^88,89^:

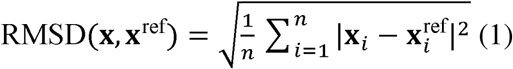

Both tools yielded similar results and we report RMSD values from the MDTraj calculations. The root-mean-square-fluctuation (RMSF) is calculated according to the equation below, where (R_i_) is the mean atomic coordinate of the *i*th C_α_ atom and R_i_ is its instantaneous coordinate:

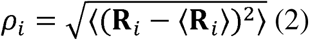

RMSF analysis and structural changes. By analyzing the trajectories of the system, we also calculate the dynamic correlation between all atoms within the molecule. This dynamic cross-correlation (DCC) between the *i*th and *j*th atoms is defined by the following equation:

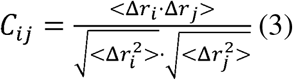

with Δ*r_i_*the displacement from the average position of atom *i*, and ⟨⟩ the time average over the whole trajectory.^90^ The DCC values are calculated between –1 and 1, where 1 corresponds to complete correlation, –1 to complete anti-correlation; and 0 indicated no correlation. The generated DCC heatmap between the *C*_α_ atoms of selected frames in a trajectory is used to identify dynamic couplings and collective motions in the protein system.

### Mutational Scanning of the RBD-Antibody Binding Interfaces

Mutational scanning analysis of the binding epitope residues for the S RBD–antibody complexes. Each binding epitope residue was systematically mutated using all substitutions, and corresponding protein stability and binding free energy changes were computed. The BeAtMuSiC approach^91–95^ was employed and evaluated the impact of mutations on both the strength of interactions at the protein–protein interface and the overall stability of the complex using statistical energy functions. BeAtMuSiC identifies a residue as part of the protein–protein interface if its solvent accessibility in the complex is at least 5% lower than its solvent accessibility in the individual protein partner(s). The binding free energy of the protein–protein complex can be expressed as the difference in the folding free energy of the complex and folding free energies of the two protein binding partners. The change in the binding energy due to a mutation was calculated then as

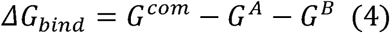

G^cum^ is the free energy of the complex. This is the Gibbs free energy associated with the folded, bound state of the entire protein–protein complex (e.g., the Spike RBD–antibody complex). G^A^ is the free energy of the first binding partner (e.g., the isolated S-RBD) in its unbound, folded state. G^B^ is the free energy of the second binding partner (e.g., the isolated antibody) in its unbound, folded state.

The change in the binding energy due to a mutation was calculated then as

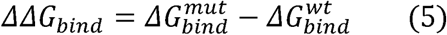

ΔΔ*G_bind_* is the change in binding free energy resulting from a specific mutation. This quantifies how the mutation affects the binding affinity compared to the wild-type (original) interaction. A positive ΔΔ*G_bind* typically indicates weakened binding (the mutation makes binding less favorable or more difficult), while a negative ΔΔ*G_bind* indicates strengthened binding (the mutation makes binding more favorable). 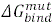 is the binding free energy calculated using Equation (4), but for the mutated protein complex (e.g., a mutant RBD bound to the antibody). 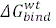 is the binding free energy calculated using Equation (4), but for the wild-type (unmutated) protein complex, serving as the reference state. We leveraged rapid calculations based on statistical potentials to compute the ensemble-averaged binding free energy changes using equilibrium samples from simulation trajectories. The binding free energy changes were obtained by averaging over 1000 and 10,000 equilibrium samples for each of the systems studied.

### Binding Free Energy Computations of the RBD Complexes with Antibodies

We calculated the ensemble-averaged changes in binding free energy using 1000 equilibrium samples obtained from simulation trajectories for each system under study. Initially, the binding free energies of the RBD–antibody complexes were assessed using the MM-GBSA approach.^96,97^ Additionally, we conducted an energy decomposition analysis to evaluate the contribution of each amino acid during the binding of the RBD to antibodies.^98,99^ The binding free energy for the RBD–antibody complex was obtained using:

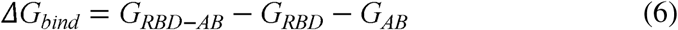

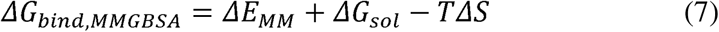

where Δ*E_MM_*is the total gas phase energy (sum of Δ*Einternal*, Δ*Eelectrostatic*, and Δ*Evdw*), and Δ*Gsol* is the sum of polar and non-polar contributions to solvation. Here, *G_RBD–ANTIBODY_* represents the average over the snapshots of a single trajectory of the complex, and *G_RBD_*and *G_ANTIBODY_* correspond to the free energy of the RBD and the antibody, respectively.

The polar and non-polar contributions to the solvation free energy were calculated using a Generalized Born solvent model and consideration of the solvent-accessible surface area.^100^ MM-GBSA was employed to predict the binding free energy and decompose the free energy contributions to the binding free energy of a protein–protein complex on a per residue basis. The binding free energy with MM-GBSA was computed by averaging the results of computations over 10,000 samples from the equilibrium ensembles. The standard error of the mean (SEM) for binding free energy estimates was calculated from the distribution of values obtained across the 10,000 snapshots sampled for each system. In this study, we chose the “single trajectory” protocol (one trajectory of the complex) because it is less noisy due to the cancellation of intermolecular energy contributions. Entropy calculations typically dominate the computational cost of MM-GBSA estimates. In this study, the entropy contribution was not included in the calculations of binding free energies of the RBD–antibody complexes because the entropic differences in estimates of relative binding affinities were expected to be small owing to the small mutational changes and the preservation of the conformational dynamics.^101,102^

### Local Frustration Analysis of Conformational Ensembles

To characterize the energetic landscape of the S-RBD complexes we performed local frustration analysis using the Frustratometer web server (http://www.frustratometer.tk).^61,62^ This approach quantifies the degree to which specific residue–residue interactions in a protein structure are energetically favorable relative to alternative states, providing insight into regions that are evolutionarily optimized (minimally frustrated), functionally adaptable (neutrally frustrated), or conformationally strained (highly frustrated). For each conformational ensemble derived from CG-CABS simulations and atomistic reconstructed ensembles we computed two types of local frustration. Mutational frustration assesses the energetic optimality of a native amino acid pair (i, j) by comparing its interaction energy to that of all viable alternative amino acid pairs at the same structural positions, while keeping the backbone geometry fixed. Configurational frustration evaluates the sensitivity of the interaction to local structural perturbations by randomizing both residue identities and interatomic distances within the native contact geometry.

In both cases, the local frustration index for a given contact was calculated as the Z-score of the native interaction energy compared to a reference ensemble of 1,000 decoys. Following established criteria from Frustratometer V2 program^103^, contacts were classified as minimally frustrated if Z > 0.78 (native interaction is significantly more favorable than alternatives), highly frustrated if Z < −1.0 (native interaction is significantly less favorable), and neutrally frustrated otherwise (native interaction is neither strongly favored nor disfavored). To assign a residue-level frustration profile, we computed the local density of frustrated contacts within a 5 Å radius of each residue.^63,64^ A residue was classified based on the dominant frustration type among its interacting partners that appeared in >50% of the structural ensemble (i.e., across the most populated conformational clusters). This approach captures both the structural stability and mutational tolerance of key functional regions—particularly the RBD–antibody interface—and links local energetic features to observed patterns of viral evolution.

## Results

### Structural Analysis of the RBD Complexes with Antibodies

We leveraged and capitalized on our earlier studies of class 1 D55-1205 antibody binding^44^ to considerably expand the scope and examine the growing set of ultrapotent and broadly neutralizing Class 1 SARS-CoV-2 neutralizing antibodies including 19-77, ZCP4C9, and ZCP3B4. Structural analysis and comparison of the binding epitopes for class 1 antibodies (Figure 2A-H, Supporting Information, Tables S2-S5) showed similar and extensive epitope coverage within the RBD. The binding epitope residues are distributed across multiple structural elements, including the receptor-binding motif (RBM), hydrophobic core, and receptor-binding ridge (Figure 2A-H). Noticeably, however, the binding epitopes for these antibodies displayed some individual peculiarities, particularly pointing to the overall denser and wider epitope featured by the most potent BD55-1205 antibody (Figure 2A, B). It is worth noting that the binding epitope of BD55-1205 spans over 20 residues, forming an extensive patch along the receptor-binding ridge, with contiguous stretches such as 454–460, 486–494, 498-505 (Figure 2A).

**Figure 2.**
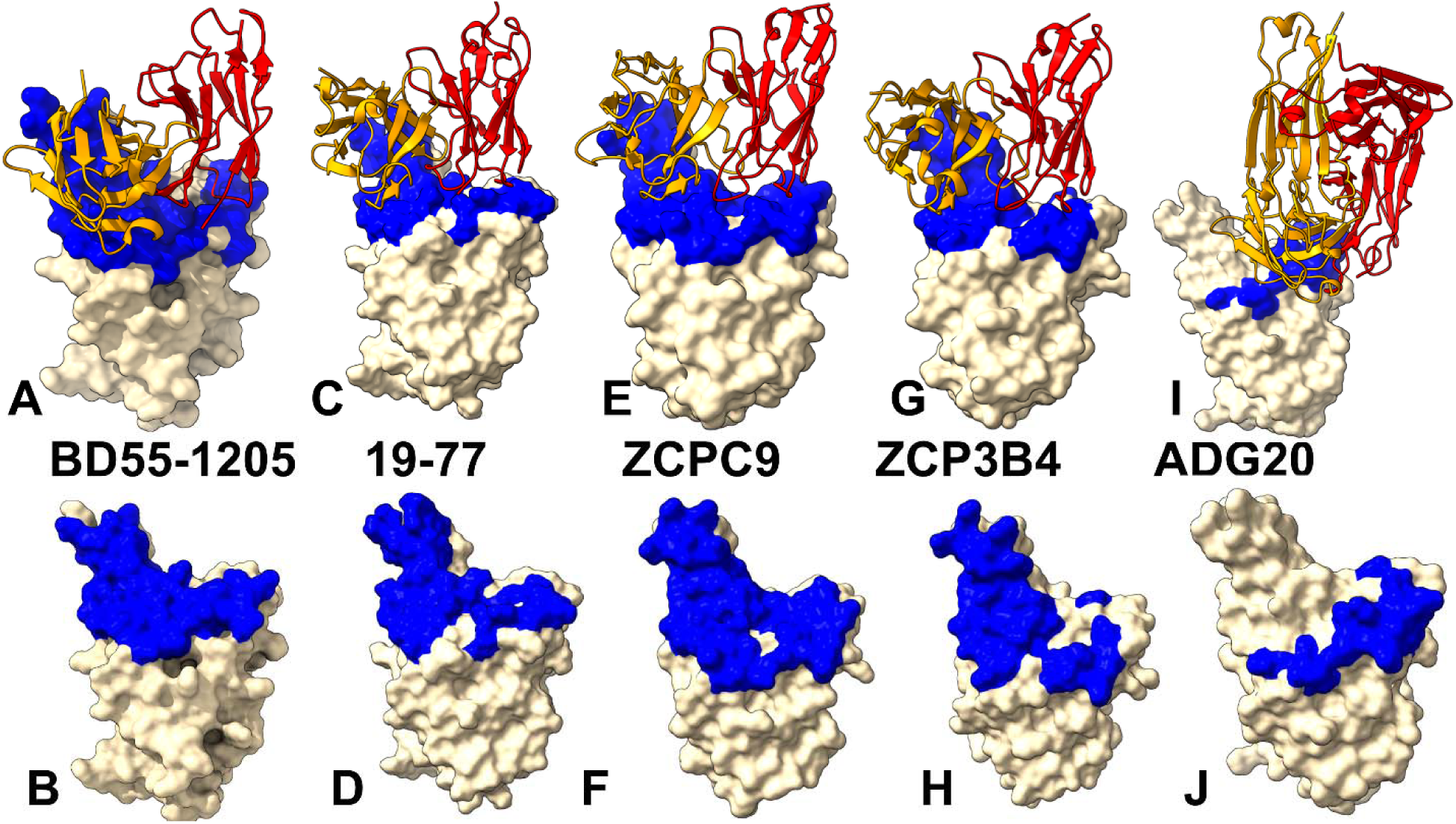
Structural organization of the RBD complexes and binding epitopes for class I antibodies. (A) The structure of class 1 BD55-1205 with XBB.1.5 RBD (pdb id 8XE9). The heavy chain in orange ribbons, the light chain in red ribbons. (B) The RBD and binding epitope footprint for D55-1205. The binding epitope residues are shown on the blue surface. (C) The structure of class 1 19-77 antibody bound with RBD (pdb id 9CFE). The heavy chain in orange ribbons, the light chain in red ribbons. (D) The RBD and binding epitope footprint for 190-77 antibody. The binding epitope residues are shown on the blue surface. (E) The structure of class 1 ZCP4C9 antibody bound to BA.5 RBD (pdb id 8K18). The heavy chain in orange ribbons, the light chain in red ribbons. (D) The RBD and binding epitope footprint for ZCP4C9. The binding epitope residues are shown on the blue surface. (G) The structure of class 1 ZCP3B4 bound with BA.5 RBD (pdb id 8K19). The heavy chain in orange ribbons, the light chain in red ribbons. (H) The RBD and binding epitope footprint for ZCP3B4. The binding epitope residues are shown on the blue surface. (I) The structure of class 4/1 ADG20 bound with RBD (pdb id 7U2D. The heavy chain in orange ribbons, the light chain in red ribbons. (H) The RBD and binding epitope footprint for ADG20. The binding epitope residues are shown on the blue surface.

A novel ultrapotent 19-77 antibody interacts essentially with the same RBD residues (403, 405, 408, 415, 416, 417, 420, 421, 455-460, 473-478, 485-490, 493, 495-502, 505) covering the entire binding interface with the ACE2 receptor (Figure 2C,D). The binding epitopes for ZCP4C9 (Figure 2E,F) and ZCP3B4 (Figure 2G, H) include contiguous stretches 403-409, 414-421, 453-460, 473-478, 486-490, 493-498, 500-505. Sequence conservation analysis revealed that the majority of this expansive epitope is relatively well-conserved across SARS-CoV-2 variants, with the notable exception of the upper half of the left shoulder of the RBD (Figure 2), which displays pronounced sequence variability—particularly at residues 455, 456, 475, and 486.^47^ Importantly, these specific residues have undergone frequent mutations in many recent subvariants EG.5.1, HK.3, JF.1, and JD.1.1 and form a subgroup of convergent evolution hotspot positions that are repeatedly exploited by the virus during evolution.

The experimentally observed exceptional neutralization profile for these class 1 antibodies^46,47^ suggest that these antibodies can accommodate mutational changes in residues 455, 456, 475, and 486. Interestingly, the scope of mutational changes featured in the very recent variants (LF.7, LF.8.1.1, NB.1.8.1, XFC, XFG) include most prominently R346T, A435S, K444R, H445R, F456L, A475V, N487D, Q493E). Apart from F456L and A475V, other RBD positions that are mutated in the latest variants do not belong to the binding epitopes of these class 1 antibodies and therefore these antibodies may be less susceptible to these changes.

A vastly different binding epitope is presented to the class 4/1 antibody ADG20 (Figure 2I, J, Supporting Information, Table S6). The binding residues include 403,405, 408,409, 439,449, 496-506. The structural analysis of the binding epitope for ADG20 showed critical role of the extended RBD segment of residues 496-506 (Figure 2I,J) making multiple contacts with Y501, G502, V503, G504, H505 and T508.^49,50^ The core of the binding epitope corresponds to the conserved RBD region (residues 436-440) that is critical for RBD stability and ACE2-binding segment 498-508 where residues are unlikely to mutate for immune escape as they are critical for ACE2 affinity.

### Conformational Dynamics of the RBD Complexes with Antibodies

We performed multiple CG-CABS simulations of the RBD-antibody complexes. The root-mean-square fluctuation (RMSF) profiles offer a detailed view of the dynamic behavior of RBD residues upon antibody binding, highlighting both shared features and notable differences among the antibodies. For reference and comparison, we also included the RMSF profiles for BD55-1205 and VIR-7229 antibodies, thus enabling a collective analysis of the RBD mobility for the entire cohort of broadly neutralizing class 1 antibodies. The RMSF analysis reveals that the core structural elements of the RBD—specifically the central β-sheet (residues 350–360), α-helices (375–380), and the loop spanning residues 394–403—exhibit low RMSF values across all class 1 antibody complexes (Figure 3A). These regions stay largely structurally conserved upon binding antibody, preserving the overall fold of the RBD. The RBD regions such as residues 355–375, which flank the RBD core and lie near the epitope interface, show moderate fluctuations, likely due to their proximity to the binding site and involvement in interfacial adjustments during complex formation (Figure 3A).

**Figure 3.**
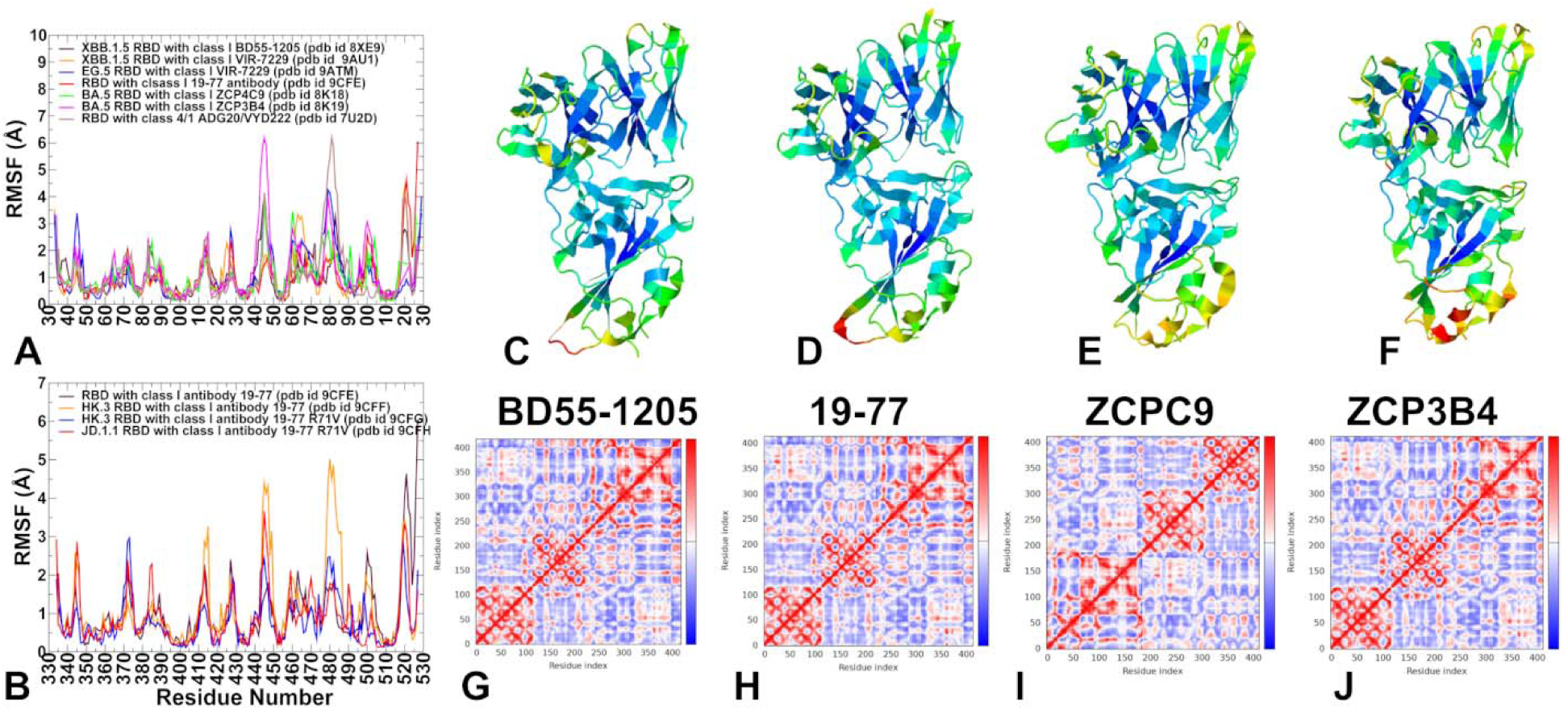
Conformational dynamics profiles obtained from CG-CABS simulations and atomistic reconstruction of the RBD-antibody complexes. (A) The RMSF profiles for the RBD residues obtained from simulations of the S-RBD complexes with class 1 antibodies: BD55-1205 with XBB.1.5 RBD, pdb id 8XE9 (in maroon lines), VIR-7229 with XBB.1.5 RBD, pdb id 9AU1 (in orange lines), VIR-7229 with EG.5 RBD, pdb id 9ATM (in blue lines), 19-77 with RBD, pdb id 9CFE (in red lines), ZCP4C9 with BA.5 RBD, pdb id 8K18 (in green lines), ZCP3B4 with BA.5 RBD, pdb id 8K19 (in magenta lines), and class 4/1 (group F3) ADG20 antibody with RBD, pdb id 7U2D (in brown lines). (B) The RMSF profiles for the RBD residues obtained from simulations of the S-RBD complexes with class 1 antibodies 19-77 with RBD, pdb 9CFE (in maroon lines), 19-77 with HK.3 RBD, pdb id 9CFF (in orange lines), 19-77 R71V with HK.3 RBD, pdb id 9CFG (in blue lines), and 19-77 R71V with JD.1.1 RBD, pdb id 9CFH (in red lines). Structural mapping of conformational mobility profiles along first three slow modes for the complex of class 1 antibodies: BD55-1205 with XBB.1.5 RBD (C), 19-77 with RBD (D), ZCP4C9 with BA.5 RBD (E) and ZCP3B4 with BA.5 RBD (F). The structures are shown in ribbons with the rigidity-to-flexibility scale colored from blue (highly rigid) to red (highly flexible). The DCC for the RBD residues in the Omicron XBB.1 RBD-ACE2 (A), Omicron XBB.1.5 RBD-ACE2 (B), Omicron BQ.1 RBD-ACE2 (C), and Omicron BQ.1.1 RBD-ACE2 complexes (D). The dynamic cross correlation (DCC) residue analysis and DCC maps for BD55-1205 with XBB.1.5 RBD (G), 19-77 with RBD (H), ZCP4C9 with BA.5 RBD (I) and ZCP3B4 with BA.5 RBD (J). Covariance matrix indicates coupling between pairs of residues, i.e. whether they experience correlated (red), uncorrelated (white) or anti-correlated (blue) motions.

These dynamics suggest a balance between keeping structural integrity and allowing subtle conformational adaptations necessary for high-affinity binding. Residues 453–460 form part of the hydrophobic core of the epitope. The RMSF values in this segment are dampened upon binding of class 1 antibodies indicating significant stabilization—a feature shared by BD55-1205, VIR-7229, 19-77 antibodies suggesting that the degree of intrinsic variability in the 455 and 456 positions is relatively moderate (Figure 3A). The interesting observation of this analysis is a particularly notable stabilization of the RBD motions in complexes with BD55-1205 and 19-77 antibodies (Figure 3A).

In these complexes, the RMSF values stay low even for generally more mobile RBD regions such as 440-450 and especially the 470-490 loop region (Figure 3A). The interfacial RBD positions involved in contacts (residues 490–505) also show reduced flexibility, consistent with their role in stabilizing the antibody-RBD interactions. According to the dynamic analysis, all class 1 antibodies employ the same heavy chain antibody residues R31, N32, Y33, P53 as BD55-1205 to form structurally conserved and yet minimal contacts with L455/F456 side-chains by avoiding deep insertion into the 455–456 pocket (Figure 3A). Instead, these antibodies make stable main-chain hydrogen bonds and van der Waals contacts with the backbone of residues 455–456 and adjacent loops (e.g., 443–450). In the complex BD55-1205 primarily engages Y449 (hydrogen bond via heavy chain), Y453 (π-stacking), L455 backbone carbonyl and F456 backbone amid. All class 1 antibodies also keep contact with the RBD backbone atoms using R31 on HCDR1 and P53 on HCDR2.

The RMSF values for the heavy and light chains of antibody residues point to very minor fluctuations of these interfacial residues (R31, N32, Y33, P53) in all class 1 antibody complexes (Supporting Information, Figure S1A-E), suggesting that this evolutionary and structurally conserved suboptimal interaction solution plays a key role in enabling tolerance to mutational changes at L455/F456 positions of the RBD. Similar to BD55-1205, other class 1 antibodies engage backbone atoms and conserved side chains critical for RBD folding and ACE2 interaction. This conservation-driven binding strategy makes it difficult for the virus to mutate without compromising fitness, resulting in a high barrier to resistance. In the 19-77 complex, the 470-490 loop region shows moderately elevated RMSF values, reflecting its inherent flexibility and functional importance (Figure 3A). Moreover, the RMSF profiles for 19-77 antibody in complexes with HK.3 RBD as well as 19-77 R71V complexes with HK.3 and JD.1.1 variants suggested a similar mobility profiles in which R71V mutation may potentially induce the increased stabilization of the flexible 470-490 regions (Figure 3B). However, our analysis of the RMSF profiles also indicated that the introduction of R71V single mutation in 19-77 does not significantly alter the RBD dynamics profiles or induce a notable increase in the mobility of the RBD interfacial residues (Figure 3B).

Structural mapping of conformational mobility profiles along first three slow modes for the complex of class 1 antibodies (Figure 3C-F) highlighted the degree of RBD stabilization induced by antibodies and also emphasized the differences between structurally rigid RBD core and moderate functional rigidity of the RBD interfaces. To examine the character of dynamic couplings and quantify correlations between motions of the RBD regions we performed the dynamic cross correlation (DCC) residue analysis. The reported DCC maps for the RBD-antibody complexes (Figure 3G-I), showing positive correlations and coupling in the RBD motions.

These results bring about a new angle to scenarios of conservation-driven or adaptability-driven binding mechanisms of antibody binding as class 1 antibodies engage the RBD through multiple suboptimal interactions across a broad stable surface, rather than relying on very few “anchor” residues. This distributed binding scenario implies that a mutation at 455 or 456 RBD positions may not necessarily result in significant conformational adaptability of the RBD and antibodies near the mutational site but rather allow for only a modest drop in the overall affinity, giving rise to other suboptimal interactions.

ADG-2 is a potent class 1/4 hybrid human monoclonal antibody that binds a highly conserved epitope in the RBD core, centered on residues 496–506 (part of the so-called “inner face” or “cryptic” epitope), which is distinct from the ACE2-binding RBM (residues 438–506) but partially overlaps its base. A generally similar picture emerged in the conformational dynamics analysis of class 4/1 antibody ADG20 binding to the RBD (Figure 3A). However, we did notice significant stabilization of the RBD residues 496-506 which is a primary RBD region targeted by ADG20 while the flexible RBM region (470-490) displayed markedly elevated level of flexibility as compared to other complexes. ADG20 participates in a network of interactions with the RBD where V503, G504 and Y505 RBD residues form contacts with the heavy chain Y58, G100, Y33, S31, Y32, F96, S97 as well as with Y31, Y91, S93, and L95 of the light chain of the antibody (Figure 3A, Supporting Information, Table S5). The RMSF values for these key interacting ADG20 residues are generally very small, suggesting that the binding interfaces are stable (Supporting Information, Figure S1F). We argue that the binding mechanism for this class may involve conservation-driven binding, where ADG20 can exploit highly conserved residues critical for viral function, stability and ACE2 binding. Although the 496–506 region and the 470–490 loop (which includes part of the RBM, notably residues 475–490) are not directly adjacent in sequence, they are spatially connected through the RBD’s β-sheet scaffold. Our results suggest that ADG-2 binding could allosterically enhance flexibility in the 470–490 loop. Indeed, when ADG-2 binds tightly to the structurally rigid 496–506 β-strand, it locks this region into a fixed conformation. The RBD is a marginally stable fold; restricting motion in one area can redistribute conformational entropy to neighboring flexible loops—particularly the 470–490 loop, which is naturally dynamic and involved in ACE2 engagement. The antibody-induced hyper mobility of the 470-490 loop may sterically or dynamically occlude access to neighboring epitopes, reducing the efficacy of cocktail antibodies. The important implication of these results is that while ADG-2’s strong binding to the conserved 496–506 RBD core confers exceptional breadth, it allosterically increases flexibility in the distal 470–490 loop—a region critical for ACE2 binding and a hotspot for immune-driven mutations. This enhanced flexibility lowers the barrier for viral escape mutations in that loop, potentially accelerating the evolution of variants resistant to other antibodies, even if ADG-2 itself stays effective. Thus, the biophysical impact of an antibody extends beyond its direct epitope, influencing the virus evolutionary landscape.

### Mutational Profiling of Antibody-RBD Binding Interfaces: Molecular Mechanisms of Immune Escape and Characterization of Binding and Escape Hotspots

Mutational scanning has enabled the identification of RBD residues that are essential both for structural stability and for high-affinity binding to the ultrapotent antibodies. Mutations that significantly destabilize the RBD–antibody complex pinpoint “hotspot” residues critical for structural integrity, high-affinity binding, and viral fitness. Across multiple ultrapotent Class 1 antibodies, including BD55-1205, 19-77, ZCP4C9, and ZCP3B4 studied here, a strikingly consistent pattern emerged. These antibodies engage the RBD through a broad, distributed interface centered on four conserved clusters: Y421/Y453/L455/F456, Y473/A475/G476, N487/Y489/Q493, and Y501/G502/H505 (Figure 4A–E). This architecture reflects a competitive inhibition strategy in which class 1 antibodies bind the “upright” RBD in direct overlap with the ACE2 interface, making extensive contacts with residues that are themselves essential for host receptor engagement—most notably Y421, Y473, and Y489. Mutations at these positions (e.g., Y421A, Y473G, Y489H) cause severe destabilization of both the antibody–RBD complex (ΔΔG > 3 kcal/mol) and the RBD fold itself, explaining their complete absence in circulating variants—the fitness cost is simply too high. In contrast, residues such as F456 and A475 represent viable escape routes. Substitutions in these positions F456L/V or A475V moderately impair antibody binding (ΔΔG ≈ 1.5–2.4 kcal/mol), and they preserve sufficient ACE2 affinity and RBD stability to be evolutionarily tolerated. Indeed, F456L and A475V have repeatedly emerged in XBB.1.5, BA.2.86, and LF.7.2.1 lineages—often in combination with L455F—to evade Class 1 neutralization.^26–33^ Notably, however, these single mutations rarely abolish activity entirely. This resilience stems from the redundant, distributed nature of Class 1 binding as the loss of interaction at F456, for example, is partially sustained by persistent contacts with neighboring residues (L455, Y473, G476) and extensive backbone-mediated hydrogen bonds involving RBD residues L455, R457, K458, Q474, A475, G476, S490, L492, and G502. Consistent with the binding studies and DMS profiling ^41,42^ and our earlier computational analysis^44^ class 1 BD55-1205 antibody stands out as a posterchild of this unique pool of broadly neutralizing antibodies. In binding of BD55-1205, mutations at F456 positions such as F456P (ΔΔG = 2.36 kcal/mol), F456V (ΔΔG = 1.86 kcal/mol), and F456L (ΔΔG = 1.54 kcal/mol) impair binding and RBD stability, though less dramatically than substitutions at Y421 or Y489 (Figure 4A). Mutational scanning of RBD residues in the complexes formed by19-77 with RBD and HK.3 RBD (Figure 4B,C) as well as complexes of 19-77 R71V mutant (Supporting Information, Figure S2) showed consistent and very similar pattern, with major binding hotspots at positions Y421, L455, F456, N487, Y489, N501/Y501 and H505.

**Figure 4.**
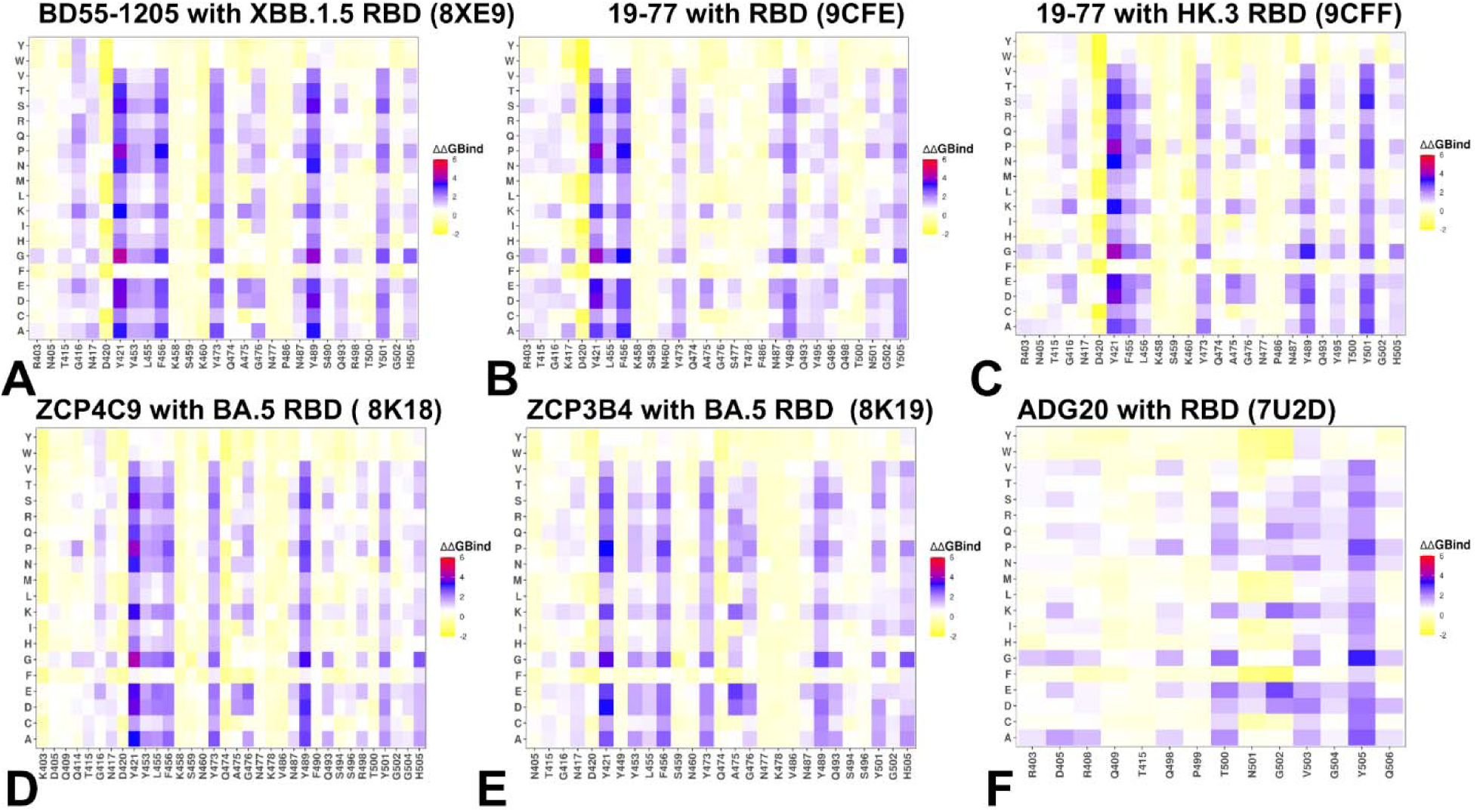
The ensemble-based mutational scanning of binding for the RBD complexes with class 1 and class 4/1 antibodies. The mutational scanning heatmaps for the binding epitope residues in the S-RBD complexes with class 1 BD55-1205 with XBB.1.5 RBD (A), 19-77 class 1 antibody with RBD (B), 19-77 class 1 antibody with HK.3 RBD (C), class 1 ZCP4C9 with BA.5 RBD (D), class 1 ZCP3B4 with BA.5 RBD (E), class 4/1 antibody ADG20 with RBD (F). The binding energy hotspots correspond to residues with high mutational sensitivity. The heatmaps show the computed binding free energy changes for 20 single mutations on the sites of variants. The squares on the heatmap are colored using a 4-colored scale blue-white-yellow-red, with blue indicating the largest unfavorable effect on binding and stability, while yellow-red points to mutations that have favorable effect and improve binding. The standard errors of the mean for binding free energy changes using randomly selected 1,000 conformational samples (0.08-0.12 kcal/mol) obtained from the atomistic trajectories.

Across all examined class 1 antibodies, the heavy chain dominates binding, with conserved hotspots at positions such as Y33, R31, P53, and L99/L100 (Supporting Information, Figures S3–S6). These residues form a flexible yet pre-configured network that engages multiple RBD segments simultaneously. For instance, in the complex with BD55-1205 antibody, Y33 contacts F456 but also contributes to interactions with Y421 and L455; similarly, L99/L100 and Y104 engage both the 455–460 loop and the 489–493 region (Supporting Information, Figures S3). This non-exclusive, multi-point anchoring implies that no single RBD mutation fully disrupts the interface. Instead, the binding energy is distributed across many moderate-affinity contacts. The light chain, by contrast, shows greater tolerance to mutation, particularly in regions contacting the conserved 500–505 patch—a segment critical for ACE2 binding and a second binding hotspot cluster for class 1 antibodies, where mutations cause destabilization (Figure 4A-E, Supporting Information, Figure S3-S6).

The results of computational mutational scanning presented here along with our previous analysis^44^ are in agreement with the DMS studies^42^ and experimental data on average antibody escape scores.^104–106^ The results identify Y473, A475, F456, G476, Y489, and L455 as top escape residues, particularly for BD55-1205 with strong agreement on the most disruptive mutations (e.g., Y473G, F456P, A475K). Overall, mutational scanning analysis for class 1 antibodies underscores a key mechanistic principle according to which this pool of antibodies can achieve ultrapotent neutralization not through a few energy-dominant “anchor” residues, but through additive, cumulative contribution of the backbone-mediated interactions across a rigid, pre-organized interface.^42^ Consistent with experimental observations^42^, this mechanism may also enable rapid on-rates and high avidity—critical for effective viral neutralization—while providing built-in redundancy against single-point escape.

The mutational scanning heatmap for ADG20 bound to the SARS-CoV-2 RBD reveals a highly structured, residue-specific pattern of energetic sensitivity that reflects the antibody unique class 4/1 binding mode — engaging both the RBM ridge (residues 403–409) and the RBM tip (residues 503–509) (Figure 4F). The ridge region (R403–R408) is a moderate-affinity anchor zone for ADG20, with D405 and R408 serving as potential moderate hotspots. Y505 showed considerable sensitivity with only Y505F showing moderate tolerance by preserving aromatic stacking while reducing side-chain bulk. This RBD region serves as a high-energy hotspot for ADG20. Class 1 antibodies typically rely heavily on residues like L455, F456, Y473, A475, which are more permissive to mutation without catastrophic fitness cost.

In contrast, ADG20 primary hotspots — R408, Y505, Q506 are less evolutionarily flexible. As a result, our results suggest that immune escape for ADG20 requires mutations at structurally or functionally constrained sites, making evasion harder and slower. Mutational scanning of ADG20 residues showed that the heavy chain energy of ADG20 extends into the 500–505 region, particularly through CDR H3 (T100) and CDR H1 (Y33). The light chain energy is centered on CDR L1 (G30/Y31/D32) and CDR L3 (Y91) that also contact T500, N501, and G502, confirming that the light chain participates in stabilizing the entire 500–505 segment (Supporting Information, Figure S7). Hence, both chains contribute to stabilizing the same set of RBD residues particularly Y505, Q506, N501, G502, and T500 (Supporting Information, Figure S7). This shared stabilization model explains why ADG20 is so resilient: even if one chain loses some affinity due to mutation, the other maintains sufficient contact to preserve overall binding. Since the 500–505 region is evolutionarily constrained, escape requires simultaneous, costly mutations.

In summary, class 1 antibodies share a conserved, evolutionarily refined binding mode: they leverage direct competition with ACE2 at functionally constrained sites while distributing binding energy across a wide interface to buffer against mutational escape. This “distributed redundancy” explains why single mutations like F456L reduce but do not abolish neutralization—and why effective immune escape typically requires combinatorial mutations across the RBM. Although it may seem somewhat paradoxical that ultrapotent antibodies would rely on relatively rigid and broadly distributed binding interfaces with suboptimal individual interactions, antibody avidity usually stems through distributed weak interactions, i.e. the additive effect of many moderate-affinity interactions across a large interface. The pre-configured, rigid interfaces minimize this penalty, enabling faster on-rates and tighter binding— especially important for neutralizing antibodies that must act quickly. As a result, mutations at 455/456 may not confer strong immune escape against Class 1 antibodies alone, explaining why these positions are not always under intense selective pressure unless combined with other mutations. It also suggests that escape requires cumulative mutations across the RBM to erode enough contacts to drop affinity below a functional threshold. The mutational scanning data confirms that ultrapotent class 1 antibodies operate through a direct competitive inhibition mechanism, leveraging strong interactions with key ACE2-contacting residues to achieve potent neutralization. ADG20, by contrast, achieves breadth through geometric and functional targeting in which cooperative binding of both heavy and light chains stabilize the same 500–505 region reducing mutational vulnerabilities, requiring cumulative mutations to fully evade binding. Unlike Class 1 antibodies which target F456 and A475 residues that are tolerable to mutations, ADG20 targets primary hotspots that are evolutionarily constrained making escape mutations rare in nature.

### MM-GBSA Analysis Reveals a Unified Blueprint for Broadly Neutralizing Antibodies: Hierarchical Energy Distribution, Hydrophobic Dominance, and Evolutionary Constraint

Through comprehensive MM-GBSA calculations across six antibody–RBD complexes — including Class 1 antibodies BD55-1205, 19-77, ZCP4C9, ZCP3B4, and the Class 4/1 antibody ADG20 — we uncover a shared energetic logic underlying broad neutralization Class 1 antibodies bind the “upright” RBD in direct competition with ACE2, engaging a broad, flat interface dominated by residues 417–490. Our MM-GBSA analysis reveals that their exceptional potency arises not from a few ultra-high-affinity anchors but rather from a hierarchical, distributed network of interactions centered on two key clusters. Residues H505, A475, Y501, Y489, N487, Y421, and L492 contribute the most favorable binding energies (ΔG ≤ –6 kcal/mol). These form a structurally conserved core cluster that overlaps extensively with the ACE2-binding site (Figure 5A-E). Their dominance is confirmed by experimental deep mutational scanning — mutations at these positions (e.g., H505Y, A475V) are among the most disruptive to BD55-1205 binding.^41,42^ L455 and F456 residues provide moderate but significant contributions, particularly through strong van der Waals interactions (F456 contributes –3.88 kcal/mol). Yet, they rank below primary hotspots in total energy placing them in a secondary tier of hotspots (Figure 5A-E). This hierarchy explains why mutations like F456L, now nearly universal in Omicron subvariants (XBB, JN.1, EG.5), cause only modest reductions in affinity rather than complete escape. The loss of interaction at F456 or L455 is buffered by the robust, synergistic network anchored by primary hotspots. This distributed, redundant architecture enables resilience without sacrificing potency. Critically, many primary hotspots including Y501, H505, Y489, N487 are also essential for ACE2 binding and RBD structural integrity. Mutations here incur high fitness costs, limiting their emergence in circulating variants. In contrast, A475V observed in JN.1 descendants modestly reduces antibody affinity while preserving ACE2 engagement, making it a viable escape route. Emerging data suggest H505Y/L may represent a rare path to partial decoupling of immune escape from fitness loss, hinting at future evolutionary pressures.^42^

**Figure 5.**
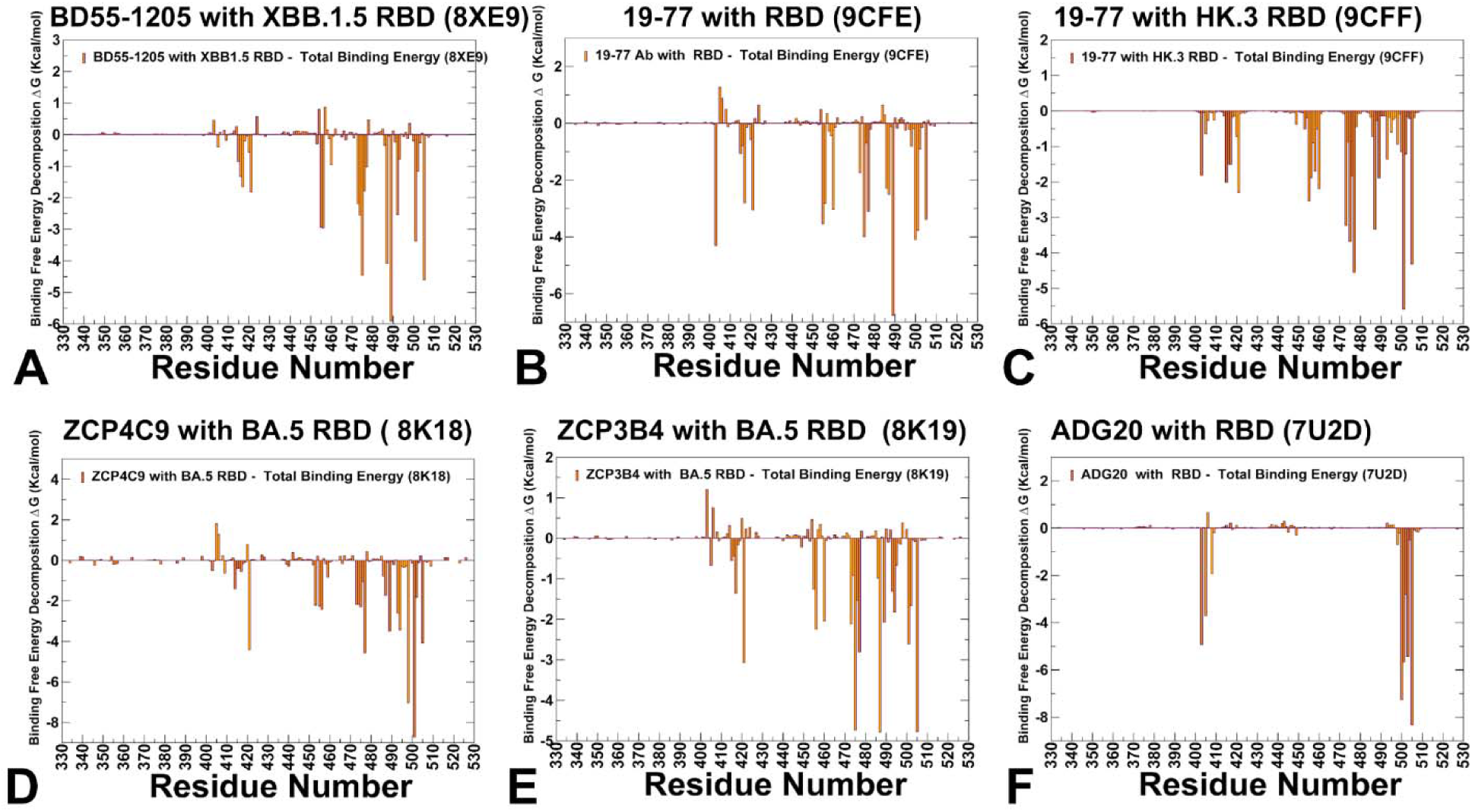
The residue-based decomposition of the binding MM-GBSA energies for the S-RBD complexes with class 1 BD55-1205 with XBB.1.5 RBD. (A), 19-77 class 1 antibody with RBD (B), 19-77 class 1 antibody with HK.3 RBD (C), class 1 ZCP4C9 with BA.5 RBD (D), class 1 ZCP3B4 with BA.5 RBD (E), class 4/1 antibody ADG20 with RBD (F). The binding free energy with MM-GBSA was computed by averaging the results of computations over 10,000 samples from the equilibrium ensembles. The standard error of the mean (SEM) for binding free energy estimates was calculated from the distribution of values obtained across the 10,000 snapshots sampled for each system. It is assumed that the entropy contributions for binding are similar and are not considered in the analysis. The statistical errors was estimated on the basis of the deviation between block average and are within 0.28-0.36 kcal/mol.

Across all class 1 antibodies the van der Waals interactions are the dominant driving force behind binding, as revealed by our residue-level VDW decomposition plots (Supporting Information, Figure S8A-E). Indeed, strong, localized van der Waals minima appear consistently at residues Y421, Y473, A475, F456, Y489, Y501, and H505 which are precisely the same residues identified as primary hotspots based on the total binding energy (Supporting Information, Figure S8A-E). This confirms that hydrophobic packing and shape complementarity are the primary determinants of affinity. The van der Waals profiles also reveal broad, shallow valleys across the interface indicating that many residues contribute modestly, reinforcing the model of additive, distributed binding. The electrostatic contributions are less significant but my contribute to buffering mutational effects (Supporting Information, Figure S9). In particular, for BD55-1205, one reason why this class 1 antibody can retain activity against variants carrying mutations F456L or L455F is that electrostatic networks of nearby residues (e.g., R457, K458, Q474) maintain favorable electrostatic interactions with the antibody and can partially compensate for loss of hydrophobic contacts (Supporting Information, Figure S9A). This synergy between VDW and electrostatics creates a robust, multi-layered interface — where loss of one interaction type is mitigated by the other (Supporting Information, Figures S8,S9). In general, class 1 antibodies exhibit a rigid, pre-organized binding mode where their interfaces are structurally complementary to the RBD where no single residue is indispensable.

This model challenges the intuitive notion that maximal potency requires deep, localized energetic hotspots. Instead, it posits that high functional affinity (avidity) arises from the additive contribution of many moderate-affinity contacts, particularly when the paratope and epitope are structurally pre-aligned. Such rigid interfaces minimize entropic penalties upon binding, enabling rapid association kinetics and tight overall binding—critical attributes for neutralizing antibodies that must intercept virions before cellular entry.

While class 4/1 ADG20 binds from a different angle, targeting the RBM without significant overlapping the ACE2 footprint, MM-GBSA profile for ADG20 reveals a striking parallel as it also relies on a hierarchical, van der Waals – dominated interface but focused on a set of functionally constrained residues. The MM-GBSA analysis showed moderate binding contributions of K403, G404, R408, while the dominant binding energy hotspots on BA.5 RBD are Y505, T500, N501 and V503 (Figure 5F). These residues form a continuous, energetically integrated surface — with both heavy and light chains contributing to stabilization of the 500– 505 loop. The van der Waals interactions dominate here as well showing deep, narrow troughs at Y505, Q506, and N501 (Supporting Information, Figure S8F). The 500–505 region has remained remarkably conserved since Omicron’s emergence in 2021 — suggesting that mutations here impose significant fitness costs. This explains why ADG20 retains broad activity despite targeting what appears to be a mutation-prone region.

This creates a high genetic barrier to resistance: escape requires mutations that simultaneously evade the antibody and preserve viral viability creating an increasingly unlikely evolutionary path. Single-point mutations (e.g., F456L, A475V) reduce affinity but rarely abolish it; true escape demands cumulative changes across multiple sites — often at significant fitness cost. Experimental DMS data confirm this: while mutations at H505 or Y501 can partially decouple immune evasion from ACE2 loss, they remain rare in circulating strains — suggesting the virus is still navigating a narrow fitness landscape.^42^ This framework may provide a predictive, design-oriented lens for evaluating therapeutic antibodies and designing next-generation vaccines.

### Conformational Frustration Profiles Reveal a Unified Strategy for Broad Neutralization in Class 1 Antibodies: Stability Through Distributed, Evolutionarily Constrained Interfaces

To examine how these balancing relationships between structural stability and plasticity we quantified conformational and mutational frustration of the RBD residues using the equilibrium conformational distributions. Conformational frustration quantifies the degree of energetic conflict a residue experiences within its local structural environment whereby neutral frustration residues represent balanced, low-energy states — often evolutionarily conserved and functionally important. Minimal frustration is a signature of highly stable structurally indispensable sites in the RBD core while high frustration may be an attribute of highly flexible, surface-exposed, or mutation-prone (Figure 6), The conformational frustration landscape of antibody–RBD complexes provides a powerful lens through which to understand why certain antibodies — particularly class 1 neutralizing antibodies such as BD55-1205, VIR-7229, 19-77, ZCP4C9, and ZCP3B4 — retain potent activity against rapidly evolving SARS-CoV-2 variants.

**Figure 6.**
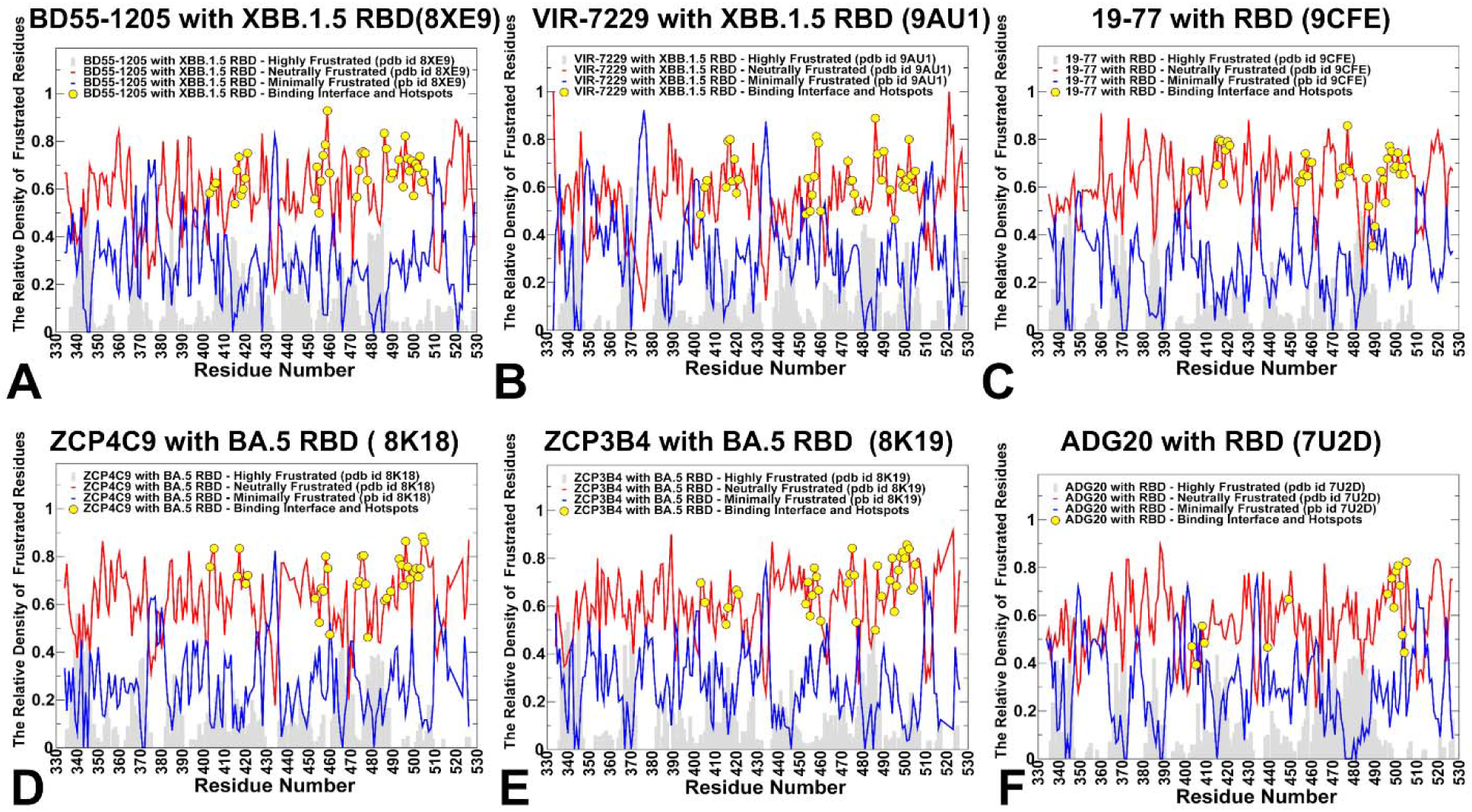
The distributions of conformational frustration for the S-RBD complexes with class 1 BD55-1205 with XBB.1.5 RBD. (A), class 1 VIR-7229 with XBB.1.5 (B), 19-77 class 1 antibody with RBD (C), class 1 ZCP4C9 with BA.5 RBD (D), class 1 ZCP3B4 with BA.5 RBD (E), and class 4/1 antibody ADG20 with RBD (F). The relative densities of high conformational frustration are shown in grey background bars, the relative densities of minimal conformational frustration are shown in blue lines and the relative densities of neutral conformational frustration are shown in red lines. The positions of the RBD binding interface residues and immune escape hotspots are highlighted in filled yellow circles.

In this analysis, we included all highly potent and broadly neutralizing class 1 antibodies that showed excellent neutralization against currently circulating new Omicron variants. In addition to 19-77, ZCP4C9, and ZCP3B4 antibodies, we included in this analysis as important baseline cases previously simulated in our studies complexes of BD55-1025 with XBB.1.5 RBD^44^ and VIR-7229 complex with XBB.1.5 RBD.^45^ Contrary to intuitive assumptions that immune escape would arise from targeting “unstable” or “highly frustrated” regions, our analysis reveals that Class 1 antibodies bind epitopes dominated by neutral frustration (Figure 6A-E), which is a signature of structural stability and evolutionary optimization and can achieve resilience by distributing binding energy across a rigid, redundant interface anchored to functionally constrained residues. Across all class 1 antibodies analyzed the RBD interface is overwhelmingly characterized by neutral conformational frustration as the RBD has evolved under strong selective pressure to maintain ACE2 binding and spike trimer stability, favoring neutral energetic landscapes. However, what distinguishes these antibodies is not the presence of neutral frustration but its spatial organization and functional context (Figure 6A-E).

Strikingly, the results revealed that binding hotspots and immune escape centers coincide with neutral frustration. Immune escape hotspots such as F456, L455, A475, Y501, H505 fall squarely within zones of neutral frustration, not high frustration. This implies that immune escape hotspots are energetically permissive to mutation exploiting neutrality of the conformational landscape to maintain the RBD; stability they simply reduce antibody affinity making them evolutionarily viable. While neutral frustration dominates, localized peaks of minimal frustration appear at residues Y421, Y489, and Y473 all of which are deeply buried or involved in core RBD structure. These residues are highly sensitive to mutation (ΔΔG > +5 kcal/mol) and are rarely mutated in circulating variants precisely because they are functionally indispensable. The revelation of this analysis suggests that neutral frustration can enable immune escape without compromising viral fitness.

We argue that the dominance of neutral frustration at escape hotspots explains a central paradox on why do mutations such as F456L or A475V persist in circulating variants. First, according to conformation frustration profiles, these residues are not under structural strain. Instead, they are neutrally frustrated and mutations in these positions do not destabilize the RBD. Second, these residues may not be directly involved in ACE2 binding or the mutation in these positions preserves sufficient ACE2 affinity. In contrast, mutations at minimally frustrated residues (e.g., Y421A, Y489H) are rare because they disrupt the RBD core fold or ACE2 interface, imposing a severe fitness cost.

Thus, neutral frustration defines the “evolutionary playground” of the virus: it marks the subset of surface residues that are both accessible to antibodies and tolerant to change. Hence. the conformational frustration profiles of Class 1 antibodies revealed that these antibodies function by binding to a stable, neutrally frustrated landscape and achieve resilience by distributing binding energy across a broad interface anchored to functionally indispensable, minimally frustrated residues. The conformational frustration profile of ADG20 is also dominated by neutral frustration across its binding interface, including the key 500–505 region. Moreover, residues Y505, Q506, N501, T500, G502, V503, and G504 all fall within zones of neutral frustration (Figure 6F). This region must remain flexible for ACE2 binding but it is functionally indispensable making mutations there evolutionarily expensive.

Our results suggest that resilience of ADG20 antibody lies in targeting neutrally frustrated and yet functionally constrained regions where escape is possible but costly. Thus, while both classes share a common biophysical framework — distributed, neutrally frustrated interfaces anchored to functionally important residues — ADG20 achieves greater resilience by targeting residues whose mutation impairs not just antibody binding, but also viral fitness.

### Mutational Frustration Profiles Reveal Evolutionary Vulnerability and Escape Pathway Predictability Across Class 1 and Class 4/1 Antibodies

While conformational frustration describes the intrinsic stability of the antibody–RBD complex, mutational frustration quantifies how mutation alters that stability revealing which residues, when mutated, relieve or worsen local strain. This provides a direct window into viral evolutionary pressure and predicts viable escape routes. Across all six complexes — BD55-1205, VIR-7229, 19-77, ZCP4C9, ZCP3B4, and ADG20 — the mutational frustration profiles are also dominated by neutral mutation frustration, reflecting that most RBD residues tolerate substitution without catastrophic destabilization. But some critical differences emerge, particularly in the density and amplitude of highly frustrated regions, the spatial distribution of minimally frustrated peaks, and the correlation between hotspots and mutation-prone sites (Figure 7). High mutation frustration flags sites where the virus can “breathe easier” after mutation making them evolutionarily favorable escape routes. For Class 1 antibodies, these sites are often peripheral allowing escape without compromising core function (Figure 7A-E). In contrast, ADG20 shows some density of high mutation frustration across RBD binding interface residues where the high mutation frustration is overwhelmed by more dominant neutral frustration in these regions (Figure 7F). The RBD regions with significant minimal mutation frustration highlights residues where mutations are highly detrimental for stability and function. In all antibodies, the minimal mutation frustration peaks appear at the RBD core residues (340–390) and ACE2-contact sites (Y421, Y473, Y489, H505) critical for receptor engagement. However, the amplitude and persistence of the minimal mutation frustration differ. For BD55-1205, these peaks are strong at Y421, Y473, Y489 confirming role of these sites as structural anchors (Figure 7A). For ADG20 minimal mutation frustration is elevated at R408, Y505, Q506 which are residues that are energetically critical for binding, even if not buried.

**Figure 7.**
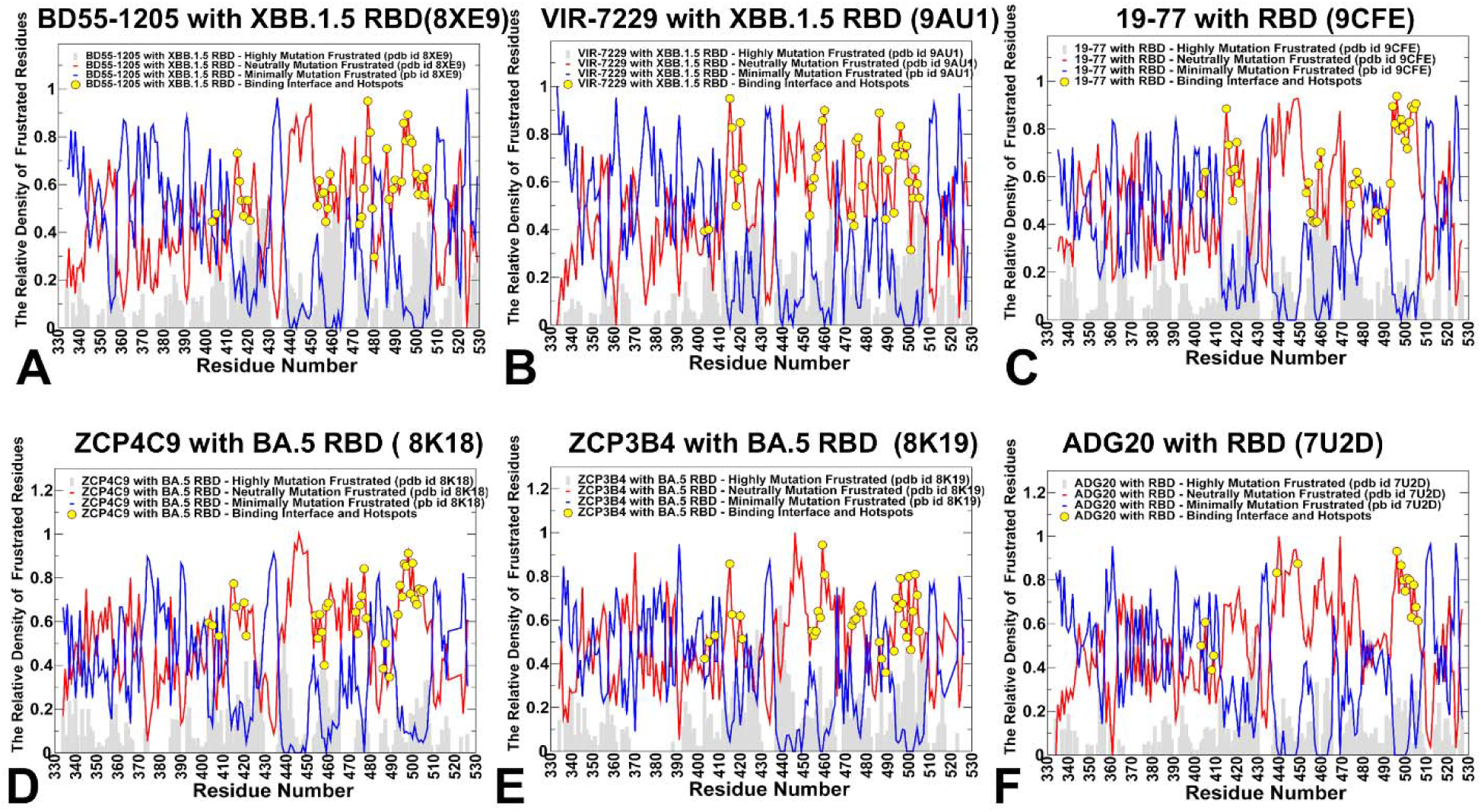
The distributions of mutational frustration for the S-RBD complexes with class 1 BD55-1205 with XBB.1.5 RBD. (A), class 1 VIR-7229 with XBB.1.5 (B), 19-77 class 1 antibody with RBD (C), class 1 ZCP4C9 with BA.5 RBD (D), class 1 ZCP3B4 with BA.5 RBD (E), and class 4/1 antibody ADG20 with RBD (F). The relative densities of high mutation frustration are shown in grey background bars, the relative densities of minimal mutation frustration are shown in blue lines and the relative densities of neutral mutation frustration are shown in red lines. The positions of the RBD binding interface residues and immune escape hotspots are highlighted in filled yellow circles.

The distributions of immune escape hotspots for BD55-1205 showed that these positions are characterized primarily by neutral mutation frustration, representing viable escape routes. Here, neutral frustration implies functional stability under mutation pressure where a neutrally frustrated residue can mutate without destabilizing the complex — and if the antibody has multiple such residues engaged, it can absorb the loss of one or two without catastrophic affinity loss (Figure 7A). Our analysis also shows that when the immune escape residue is neutrally frustrated, the virus can mutate it repeatedly — as seen with F456L/V/S/P in XBB, JN.1, EG.5. At the same time, escape is often incomplete as the antibody retains partial activity. As a result, functionally relevant immune evasion usually requires cumulative mutations across multiple hotspots (e.g., F456L + A475V).

Although the dominant neutral mutation frustration trend is generally preserved across class 1 antibodies, some immune escape positions may feature an appreciable density of minimal frustration (Figure 7B-E). When an immune escape hotspot shows appreciable density of both neutral and minimal mutation frustration, this may imply a structurally stable yet functionally critical position where mutations are tolerated but costly. Antibodies that target such dual-frustration hotspots (e.g., ADG20 at Y505/Q506, or Class 1 antibodies at Y489/H505) achieve strong binding because the residue is in a stable, well-defined conformation (minimal frustration), yet the interface remains evolutionarily relevant because the site is surface-exposed and mutable (neutral frustration). When a hotspot has both signatures, single mutations (e.g., Y505F, H505Y) may reduce antibody binding but also impair ACE2 affinity or RBD stability. As a result, such mutations emerge slowly, often only under intense immune pressure and in combination with compensatory changes. This explains why ADG20 retains activity against most Omicron subvariants—despite targeting Y505/Q506—because escape mutations here are fitness-reducing. For antibodies such as ADG20 or BD55-1205, engaging such sites is not a vulnerability—it is a deliberate exploitation of viral evolutionary constraints. The virus can mutate these residues, but doing so risks its own functionality—creating a high genetic barrier to resistance. This pattern confirms that dual-frustration hotspots may act as evolutionary “speed bumps” by lowing, but not stopping, escape.

Structural mapping of neutral frustration density—defined as residues with both conformational and mutational neutral frustration indexes exceeding a threshold of 0.7—reveals distinct yet functionally significant patterns for the Class 1 antibody BD55-1205 (Figure 8A–C) and the Class 4/1 hybrid antibody ADG20 (Figure 8D–F). In the BD55-1205–RBD complex, neutral frustration is broadly distributed across the RBD, extending well beyond the immediate binding interface to encompass functional regions frequently targeted by Omicron subvariants. Notably, the vast majority of the epitope, including key RBM segments and established binding and immune escape hotspots, is dominated by neutrally frustrated residues. This widespread neutral frustration creates an evolutionarily permissive landscape that allows for mutational exploration without catastrophic loss of structure or function. Embedded within this flexible periphery is a compact “central island” of minimally frustrated residues (403, 404, 416–420, 452, 453), which exhibit high energetic stability and structural rigidity. This minimally frustrated core likely serves as a stable anchoring point for BD55-1205, while the surrounding sea of neutral frustration provides adaptability. Together, this architecture enables a distributed, redundant binding mode that maintains high potency while conferring resilience to mutation-driven escape.

**Figure 8.**
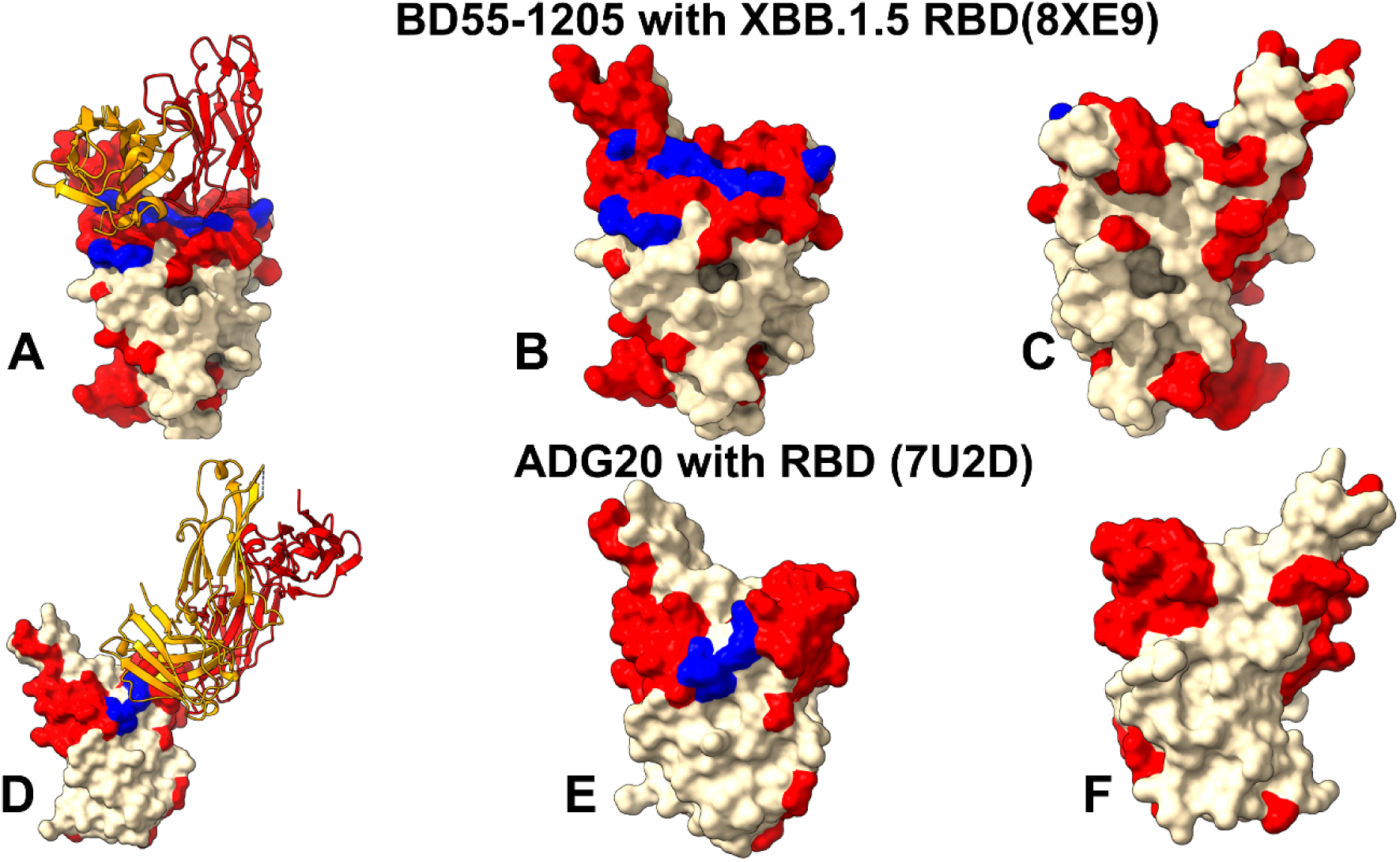
Structural mapping of neutral frustration density in complexes of class 1 antibody BD55-1205. (A-C) and class 4/1 ADG20 (D-F). The mapped neutrally frustrated residues with both conformational and mutational neutral frustration indexes exceeding a threshold of 0.7 are shown in red surface. The binding epitope residues that do not show dominant neutral frustration but are characterized by strong minimal frustration are shown in blue surface. The heavy chain in orange ribbons, the light chain in red ribbons on panels (A) and (D).

In contrast, the ADG20–RBD complex displays a more localized but equally strategic frustration pattern. Its epitope includes a small, minimally frustrated segment (residues 403–409) that may function as a structurally stable anchor. However, the functionally critical RBM tip (residues 503–508)—which dominates ADG20 binding affinity and governs its immune escape profile—is characterized by dominant neutral frustration (Figure 8D-F). This duality reflects ADG20 unique binding mechanism: it combines a rigid, conserved anchor with a dynamically adaptable, neutrally frustrated interface at the RBM apex. This configuration allows ADG20 to maintain strong binding to evolutionarily constrained regions while accommodating subtle conformational or mutational changes in adjacent residues thereby balancing potency with evolutionary robustness. Collectively, these frustration landscapes illustrate how different antibody classes exploit distinct energetic architectures to achieve durable neutralization: BD55-1205 through a broad, redundant interface enriched in neutral frustration, and ADG20 through a bipartite strategy that merges minimal frustration anchoring with neutral frustration–mediated adaptability.

A central insight from mutational frustration analysis is that immune escape hotspots overwhelmingly coincide with neutrally frustrated residues—not because these sites are unstable or disordered, but precisely because they are energetically permissive to mutation. This observation resolves an apparent paradox: the most frequently mutated residues in circulating SARS-CoV-2 variants are not located in highly frustrated (structurally strained) regions, but rather in zones of neutral frustration, where the local energetic environment is balanced and tolerant of change. In such regions, mutations do not significantly perturb the protein’s folding stability or functional conformation. From the virus’s perspective, this is ideal: it can alter surface-exposed residues to evade antibody recognition without paying a fitness cost in terms of RBD misfolding, reduced ACE2 affinity. Critically, broad, distributed interface in class I antibodies can withstand single mutations through binding redundancy. Similarly, class 4/1 ADG20, while also targeting neutrally frustrated residues, gain resilience by engaging multiple subregions (e.g., 403–409 and 503–509) where mutations may carry higher fitness costs due to indirect structural roles (e.g., maintaining RBD “up” conformation). The convergence of these observations — the universal dominance of neutral frustration, the localized significance of minimally frustrated residues, and the marginal presence of highly frustrated background — paints a coherent picture.

Our results suggest that the emerging dominant pattern of neutral frustration in the evolutionary hotspots on the RBD may create a state of “energetically suboptimal, adaptable frustration” that limits access to potentially superior adaptive outcomes but generates mutational “hotspots” – genomic regions where mutations repeatedly and predictably occur, leading to highly repeatable evolutionary trajectories. The discovered relationships indicate that neutral frustration at the key adaptable evolutionary hotspots of the RBD-antibody interfaces can create mutational path leading to rapid adaptation of immune escape mutations that represent suboptimal and yet robust recurrent outcomes. Adaptable evolutionary sites contain a mixture of neutral and low frustration to balance stability and functional flexibility.

## Discussion

Our integrated analysis of structural epitopes, conformational dynamics, binding energetics, and frustration landscapes across multiple ultrapotent Class 1 (BD55-1205, 19-77, ZCP4C9, ZCP3B4) and Class 4/1 (ADG20) antibodies reveals a shared biophysical blueprint for broad neutralization—one that reconciles apparent contradictions between high potency, mutational resilience, and evolutionary vulnerability. Class 1 neutralizing antibodies bind via rigid, pre-configured interfaces that distribute binding energy across a broad epitope through numerous suboptimal, yet synergistic, interactions many of which occur at sites of neutral local frustration. This architecture reconciles an apparent paradox: how can ultrapotent antibodies rely on “suboptimal” contacts and still neutralize effectively? The answer lies not in maximizing affinity at a few residues, but in improving robustness through redundancy, backbone engagement, and strategic targeting of evolutionarily constrained regions. This “distributed redundancy” model explains why class 1 antibodies retain activity against variants carrying mutations at key sites like F456L or A475V—loss of interaction at one residue is buffered by persistent contacts with neighboring backbone atoms and side chains (e.g., L455, Y473, G476, Q474). Critically, these antibodies engage the RBD through extensive backbone-mediated hydrogen bonds and van der Waals packing, minimizing entropic penalties and enabling rapid on-rates—essential for intercepting virions before cellular entry. This architecture is likely an evolutionarily refined solution. By avoiding deep insertion into mutable pockets (e.g., the 455–456 cleft) and instead forming suboptimal yet stable contacts with conserved backbone elements, these antibodies achieve a balance between specificity and adaptability. The result is an interface that is structurally rigid yet functionally robust—capable of absorbing single-point mutations without catastrophic loss of affinity. Mutational scanning and MM-GBSA analyses further refine this model by revealing a hierarchical organization of binding energy. Primary hotspots such as H505, Y501, Y489, and Y421—contribute disproportionately to binding affinity and overlap directly with ACE2-contact sites. Mutations here incur severe fitness costs, explaining their near-complete absence in circulating variants. In contrast, secondary hotspots like F456 and L455 provide moderate energetic contributions but reside in peripheral, mutation-tolerant regions— making them frequent targets of immune escape. This hierarchy creates a high genetic barrier to resistance: effective evasion requires cumulative mutations across multiple sites—often combining changes in secondary hotspots (e.g., F456L + A475V) with rare, fitness-tolerant substitutions in primary regions (e.g., H505Y). The virus thus navigates a narrow fitness landscape, where immune escape is possible but evolutionarily expensive.

The interesting insights come from conformational and mutational frustration analyses. Both class 1 and Class 4/1 antibodies bind epitopes dominated by neutral frustration—a signature not of instability, but of evolutionary optimization. Immune escape hotspots (e.g., F456, A475, Y505) fall precisely within these neutrally frustrated zones, confirming that escape arises not from targeting unstable regions, but from exploiting permissive ones. Within this shared framework, local frustration analysis provides a critical new lens: the majority of key binding hotspots and immune escape positions—including L455, F456, A475, Y473, and Y489—are characterized by neutral frustration. Neutral frustration indicates that these residues occupy energetically suboptimal but evolutionarily permissible states—neither strongly stabilizing (low frustration) nor destabilizing (high frustration)—thereby permitting mutational exploration without catastrophic loss of protein fold or function. This “energetically permissive” landscape explains why mutations at these sites recur across variants: they represent accessible, low-cost paths to immune escape that preserve viral fitness. Local frustration analysis further clarifies why certain regions become evolutionary hotspots. Neutral frustration at L455/F456 creates a mutationally permissive zone—a “neutral network” that allows genetic drift without functional loss, generating a reservoir of diversity that can be rapidly selected under immune pressure. This is consistent with the theory that neutral mutations enable access to adaptive pathways: a mutation like F456L may be nearly neutral in isolation but becomes advantageous in the context of other immune pressures. Meanwhile, low-frustration regions (e.g., the RBD core around Y421/Y489) maintain structural integrity, ensuring that escape mutations remain constrained to the periphery.

In summary, we propose a generalized model for durable Class 1 neutralization: ultrapotent antibodies achieve resilience by engaging a broad, rigid interface enriched in neutral-frustration sites, leveraging backbone contacts and energetic redundancy to buffer against mutations that would otherwise drive escape. This design exploits the virus evolutionary constraints—targeting functionally indispensable residues while tolerating variation at permissive, neutral-frustration hotspots. Moving forward, efforts to elicit or engineer antibodies with this architecture— emphasizing distributed binding, backbone engagement, and compatibility with neutral-frustration landscapes—may yield more robust countermeasures against not only SARS-CoV-2 but also future pandemic threats.

## Conclusion

The ongoing evolution of SARS-CoV-2 variants suggests that at the molecular level, the evolution of immune escape hotspots in SARS-CoV-2 is influenced through complex an nuanced confluence of genetic mutations and the diversity of antibody responses. In this study, we combine structural, dynamic, energetic, and frustration-based analyses of the expanded set of ultrapotent and broadly neutralizing Class 1 SARS-CoV-2 neutralizing antibodies—BD55-1205, 19-77, ZCP4C9, and ZCP3B4 and class 4/1 ADG20 antibody to reveal a unifying biophysical principle that explains how these antibodies achieve both exceptional potency and remarkable resilience against viral evolution. Using multi-pronged computational approach, we found these neutralizing antibodies bind via rigid, pre-configured interfaces that distribute binding energy across a broad epitope through numerous suboptimal, yet synergistic, interactions—many of which occur at sites of neutral local frustration. While the growing pool of broadly neutralizing class I antibodies often share a general binding mode, these studies highlighted that subtle differences in atomic-level interactions can critically shape each antibody unique neutralization profile. The results of this study indicate that the distributed, rigid, and frustration-neutral binding mode for class 1 antibodies can establish a high genetic barrier to resistance as immune escape would require the cumulative erosion of binding energy through multiple mutations.

Dynamically, these antibodies impose significant stabilization on the RBD, suppressing fluctuations not only at direct contact sites (e.g., residues 453–460 and 490–505) but also in distal loops (e.g., 470–490), thereby locking the RBD into a conformation that sterically occludes ACE2 binding. This rigidity minimizes entropic penalties upon association, enabling rapid on-rates and tight complex formation—critical for effective viral neutralization. Importantly, this lack of conformational adaptability means the system does not “rescue” binding through induced fit when mutations occur; instead, it absorbs perturbations through energetic redundancy, resulting in only modest affinity losses. Energetically, MM-GBSA decomposition reveals a two-tiered hotspot architecture: a primary tier (H505, A475, Y501, Y489, Y421) that dominates binding free energy and overlaps with ACE2-contact residues, and a secondary tier (L455, F456, G476, Y473) that contributes auxiliary—but non-essential—interactions. This hierarchy explains why mutations like F456L or A475V, now nearly fixed in XBB, JN.1, and LF lineages, reduce but do not abolish neutralization: the loss at secondary sites is buffered by the robust network anchored in evolutionarily constrained primary residues

Crucially, local frustration analysis links this biophysical architecture to viral evolutionary mechanisms. We find that the most often mutated escape positions—L455, F456, A475—reside in regions of neutral frustration, where the energetic landscape is neither stabilizing nor destabilizing. This neutrality creates a mutationally permissive zone: mutations can accumulate with minimal impact on RBD folding or ACE2 affinity, enabling the virus to explore immune escape without severe fitness costs. In contrast, primary hot spots like Y489 and H505 exhibit low frustration and are embedded in structurally or functionally essential regions, rendering them evolutionarily constrained. Thus, the interplay between neutral frustration (enabling variation) and low frustration (enforcing conservation) shapes the virus’s adaptive landscape, directing evolution toward predictable, convergent mutations at the periphery of the epitope. This framework also explains the emergence of recent variants: mutations such as F456L, A475V, and L455F are not random—they are recurrent solutions selected from a limited set of viable escape routes permitted by the neutral-frustration architecture of the RBM. The virus navigates a narrow fitness corridor, where immune evasion must be balanced against functional viability. Antibodies that exploit this constraint—by targeting conserved, low-frustration cores while tolerating variation at neutral-frustration peripheries—create a high barrier to resistance.

This highlights that antibody efficacy must be evaluated not only by direct neutralization potency but also by its systemic impact on viral evolvability. Together, our findings suggest that durability of broadly neutralizing antibodies may arise from optimal energetic distribution across a rigid, backbone-engaging interface embedded within a frustration landscape that mirrors the virus own evolutionary constraints.

## Author Contributions

Conceptualization, G.V.; methodology, G.V.; software, M.A., V.P., B.F., G.H., G.V.; validation, G.V.; formal analysis, G.V., M.A., G.H., investigation, G.V.; resources, G.V., M.A. and G.H.; data curation, G.V.; writing—original draft preparation, G.V.; writing—review and editing, G.V., M.A. and G.H.; visualization, V.P., B.F., G.V.; supervision, G.V.; project administration, G.V.; funding acquisition, G.V. All authors have read and agreed to the published version of the manuscript.

## Data Availability Statement

Data is fully contained within the article and Supplementary Materials. Crystal structures were obtained and downloaded from the Protein Data Bank (http://www.rcsb.org). The rendering of protein structures was done with UCSF ChimeraX package (https://www.rbvi.ucsf.edu/chimerax/) and Pymol (https://pymol.org/2/). All mutational heatmaps were produced using the developed software that is freely available at https://alshahrani.shinyapps.io/HeatMapViewerApp/.

## Acknowledgments

The authors acknowledge support from Schmid College of Science and Technology at Chapman University for providing computing resources at the Keck Center for Science and Engineering.

## For Table of Contents Use Only

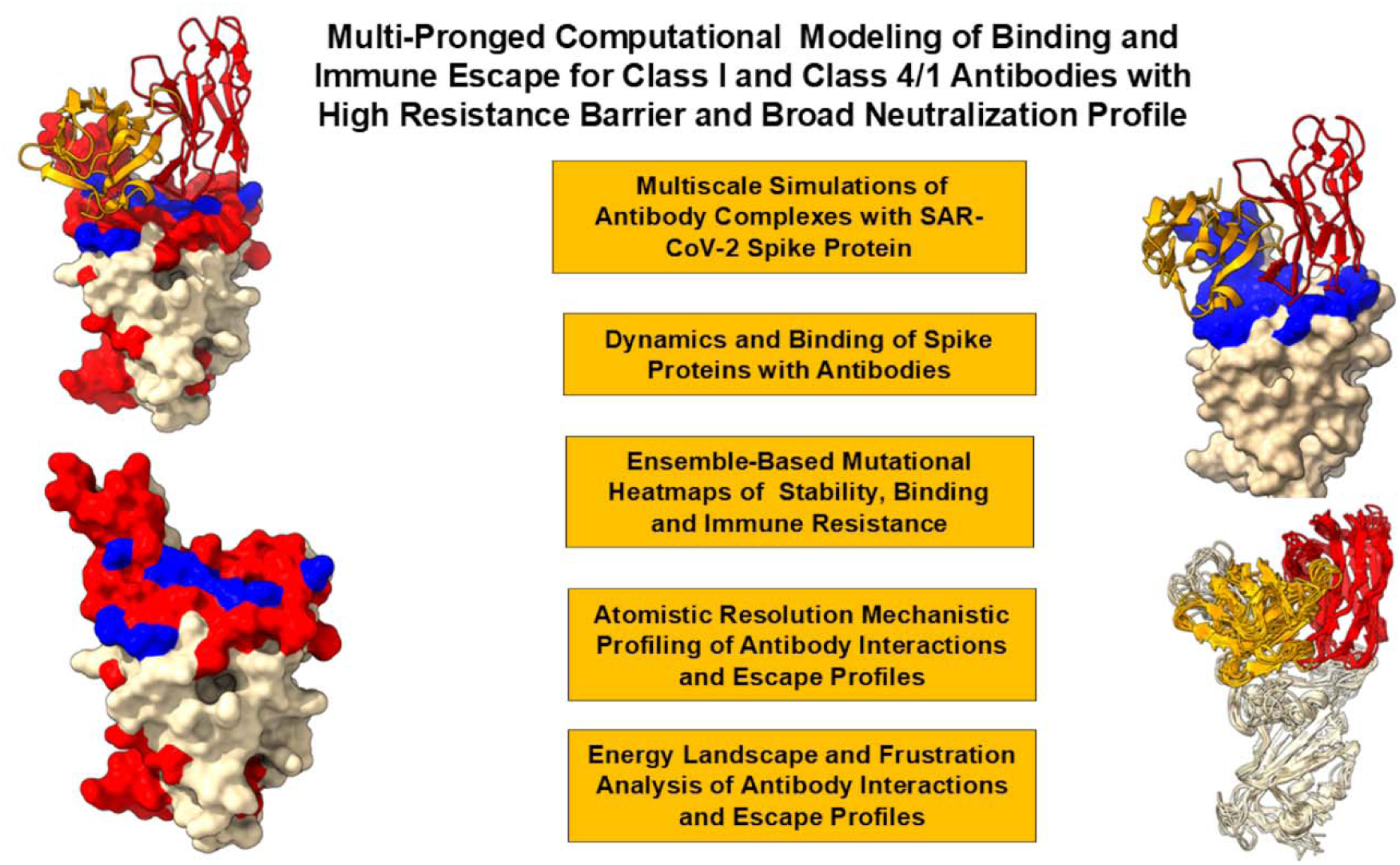

