## Supplemental Figures S1-S9 and Tables S1-S6 for "Neutral Frustration Landscape Architecture of SARS-CoV-2 Spike-Antibody Interfaces Shapes Immune Evasion Mechanisms for Ultrapotent Neutralizing Antibodies and Determines Pathways of Viral Adaptation: Insights from Integrative Computational Approach"

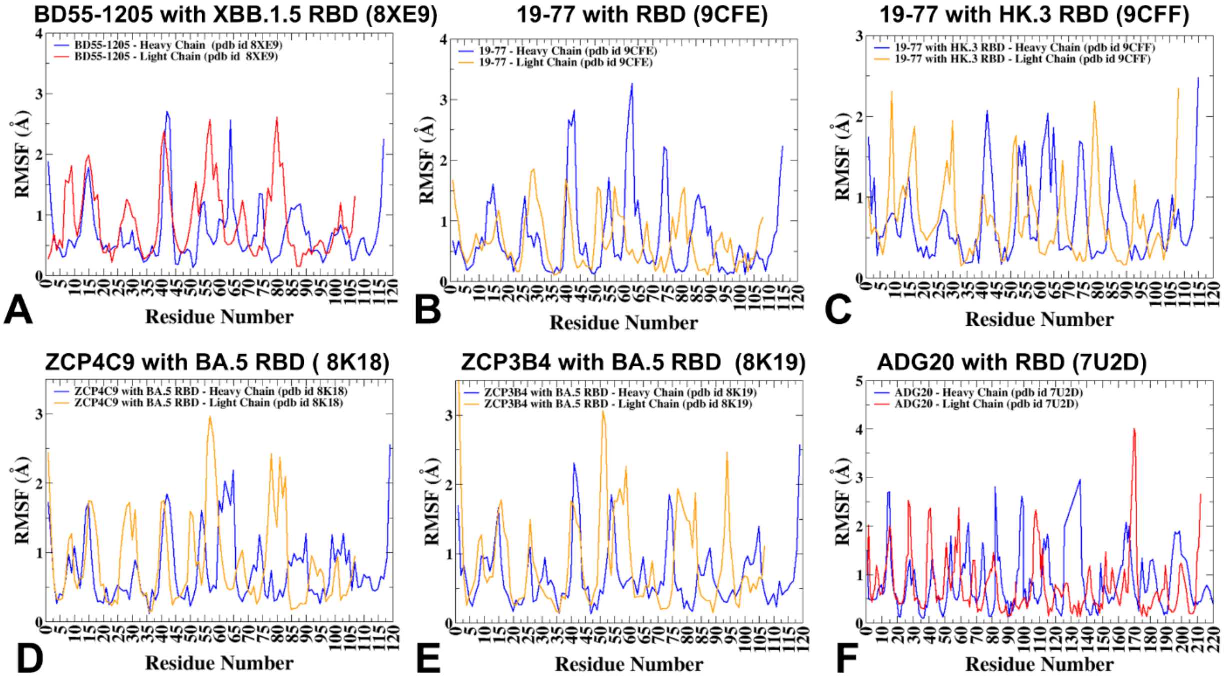


**Figure S1. Conformational dynamics profiles obtained from CG-CABS simulations and atomistic reconstruction of the RBD-antibody complexes**. The RMSF profiles for the heavy chain residues ( in bule lines) and light chain residues (in orange lines) for BD5-1205 obtained from simulations of the S-RBD complexes with BD55-1205, pdb id 8XE9 (A), for 19-77 heavy and light chains with RBD, pdb id 9CFE (B), for 19-77 heavy and light chains with HK.3 RBD, pdb id 9CFF (C), for ZCP4C9 heavy and light chains with BA.5 RBD, pdb id 8K18 (D), for heavy and light chains of ZCP3B4 with BA.5 RBD, pdb id 8K19 (E), and for heavy and light chains of class 4/1 (group F3) ADG20 antibody with RBD , pdb id 7U2D (F).

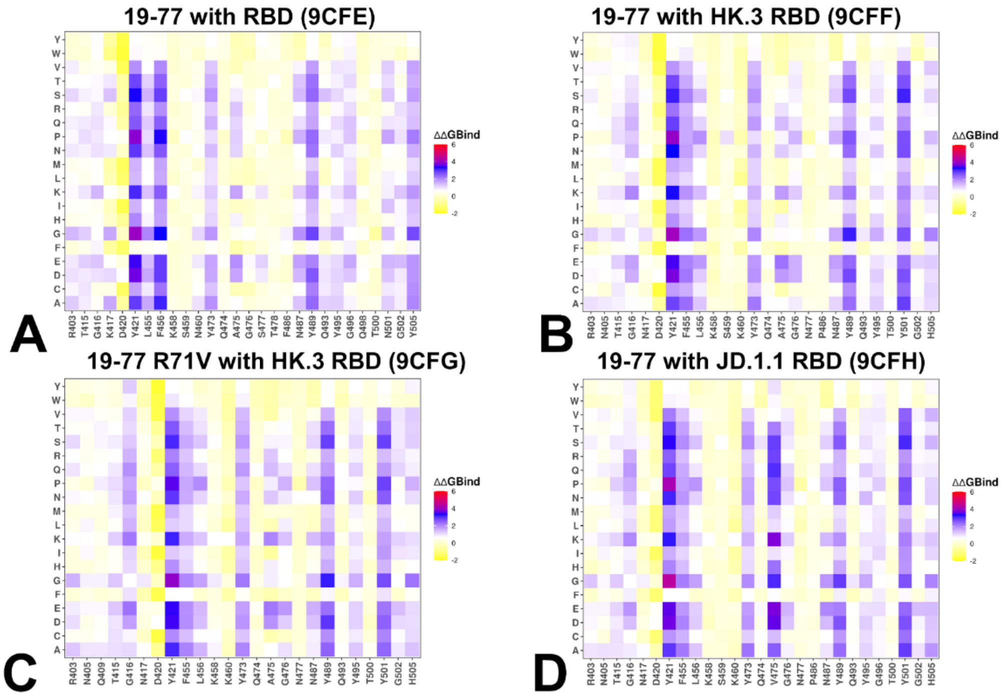


**Figure S2. The ensemble-based mutational scanning of binding for the RBD complexes with class 1 19-77 antibody.** The mutational scanning heatmaps for the binding epitope residues for 19-77 class 1 antibody with RBD, pdb id 9CFE (A), 19-77 class 1 antibody with HK.3 RBD, pdb id 9CFF (B), 19-77 R71V class 1 antibody with HK.3 RBD, pdb id 9CFG (C), and 19-77 R71V class 1 antibody with JD.1.1 RB, pdb id 9CFH (D). The binding energy hotspots correspond to residues with high mutational sensitivity. The heatmaps show the computed binding free energy changes for 20 single mutations on the sites of variants. The squares on the heatmap are colored using a 4-colored scale blue-white-yellow-red, with blue indicating the largest unfavorable effect on binding and stability, while yellow-red points to mutations that have favorable effect and improve binding.

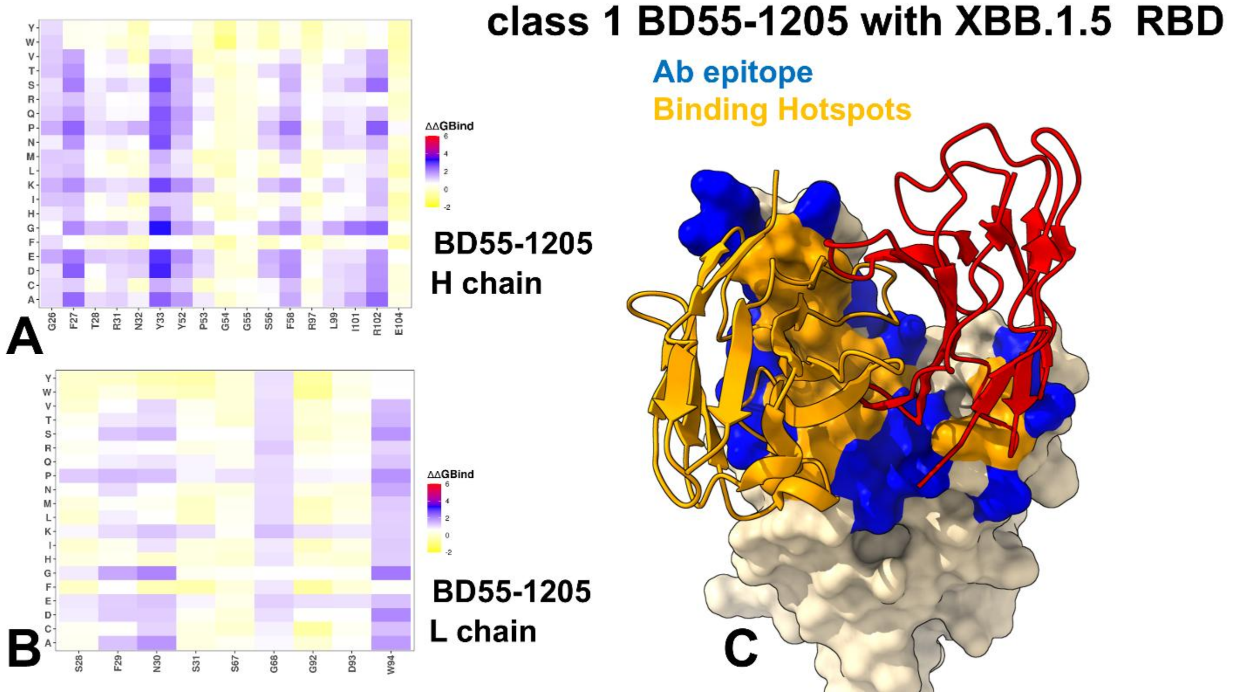


**Figure S3. Ensemble-based dynamic mutational profiling of the RBD intermolecular interfaces in the RBD complex with BD55-1205 antibody**. The mutational scanning heatmaps are shown for the interfacial heavy chain residues of BD55-1205 (A) and light chain residues of BD55-1205 (B). The heatmaps show the computed binding free energy changes for 20 single mutations of the interfacial positions. (C) The structure of BD55-1205 bound to RBD. The heavy chain of BD55-1205 is in orange ribbons, and light chain is in red ribbons. The binding epitope is shown in blue surface and the positions of the RBD binding energy hotspots are shown in orange-colored surface. (D). RBD from the complex with BD55-1205. The binding epitope residues are in blue surface and the binding interfacial RBD hotspots are in orange surface.

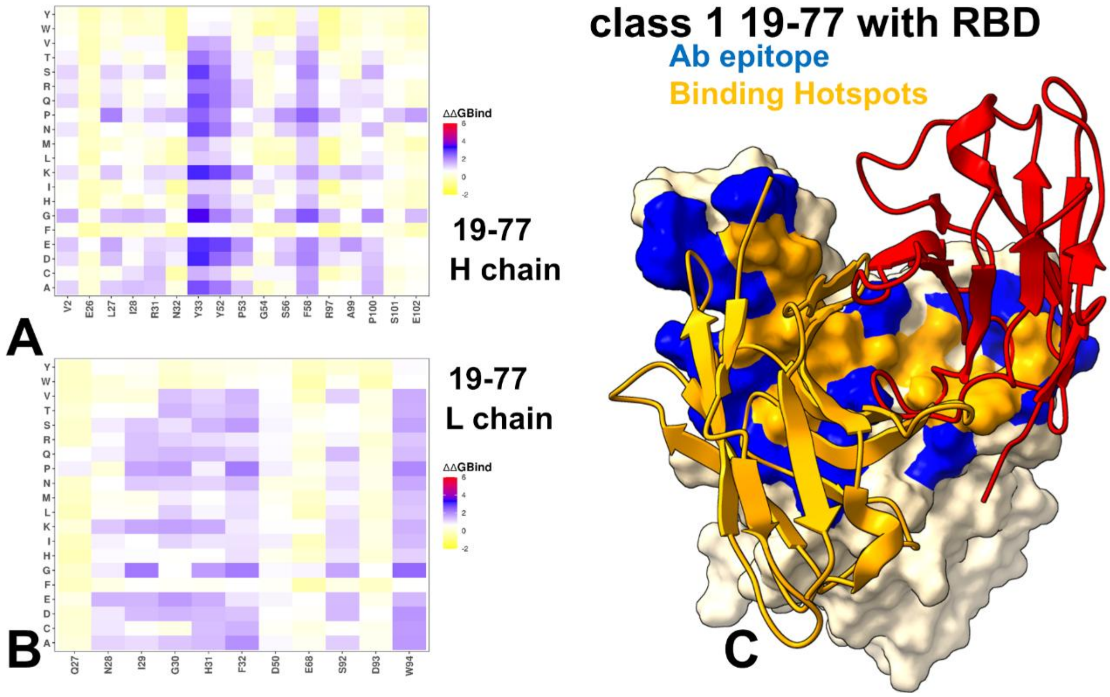


**Figure S4. Ensemble-based dynamic mutational profiling of the RBD intermolecular interfaces in the RBD complex with 19-77 antibody.** The mutational scanning heatmaps are shown for the interfacial heavy chain residues of 19-77 (A) and light chain residues of 19-77 (B). The heatmaps show the computed binding free energy changes for 20 single mutations of the interfacial positions. (C) The structure of 19-77 bound to RBD. The heavy chain of 19-77 is in orange ribbons, and light chain is in red ribbons. The binding epitope is shown in blue surface and the positions of the RBD binding energy hotspots are shown in orange-colored surface. (D). RBD from the complex with 19-77. The binding epitope residues are in blue surface and the binding interfacial RBD hotspots are in orange surface.

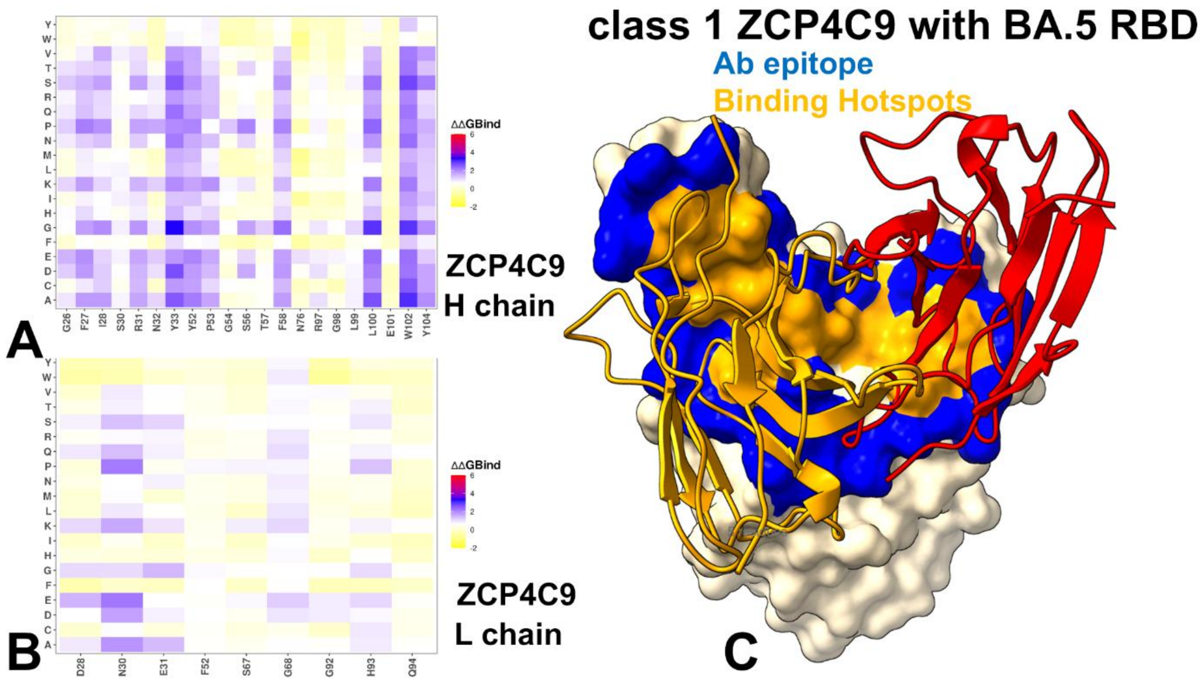


**Figure S5. Ensemble-based dynamic mutational profiling of the RBD intermolecular interfaces in the RBD complex with ZCP4C9 antibody.** The mutational scanning heatmaps are shown for the interfacial heavy chain residues of ZCP4C9 (A) and light chain residues of

ZCP4C9 (B). The heatmaps show the computed binding free energy changes for 20 single mutations of the interfacial positions. (C) The structure of ZCP4C9 bound to RBD. The heavy chain of ZCP4C9 is in orange ribbons, and light chain is in red ribbons. The binding epitope is shown in blue surface and the positions of the RBD binding energy hotspots are shown in orange-colored surface. (D). RBD from the complex with ZCP4C9. The binding epitope residues are in blue surface and the binding interfacial RBD hotspots are in orange surface.

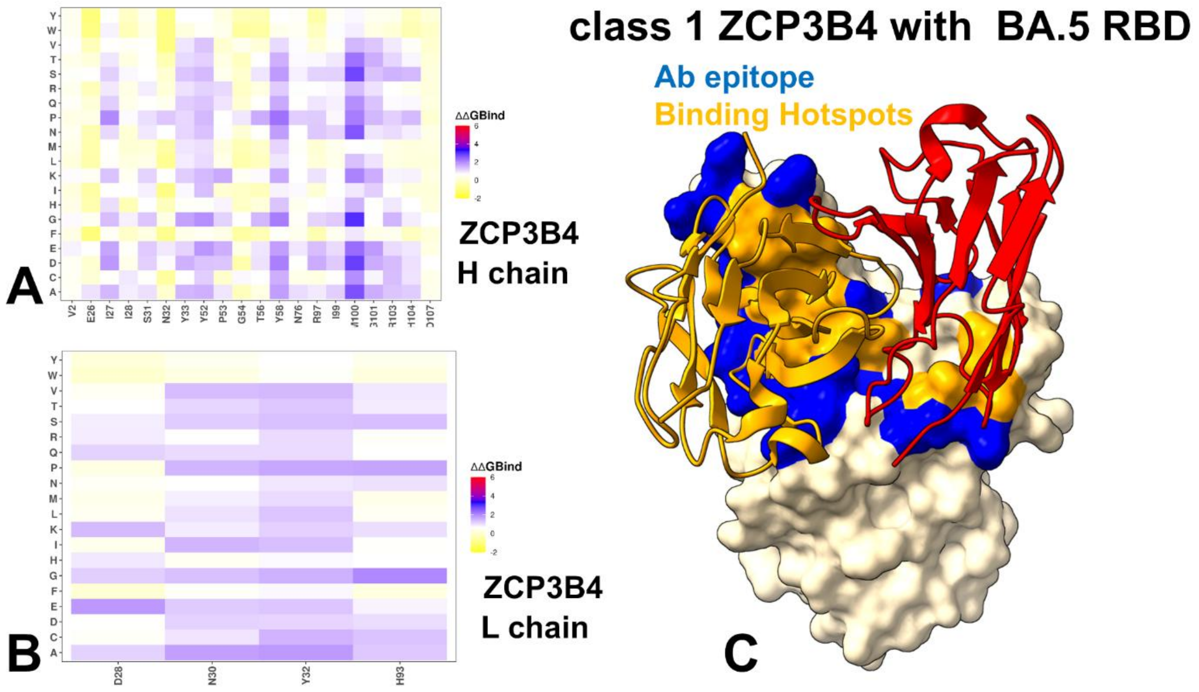


**Figure S6. Ensemble-based dynamic mutational profiling of the RBD intermolecular interfaces in the RBD complex with ZCP3B4 antibody.** The mutational scanning heatmaps are shown for the interfacial heavy chain residues of ZCP3B4 (A) and light chain residues of

ZCP3B4 (B). The heatmaps show the computed binding free energy changes for 20 single mutations of the interfacial positions. (C) The structure of ZCP3B4 bound to RBD. The heavy chain of ZCP3B4 is in orange ribbons, and light chain is in red ribbons. The binding epitope is shown in blue surface and the positions of the RBD binding energy hotspots are shown in orange-colored surface. (D). RBD from the complex with ZCP3B4. The binding epitope residues are in blue surface and the binding interfacial RBD hotspots are in orange surface.

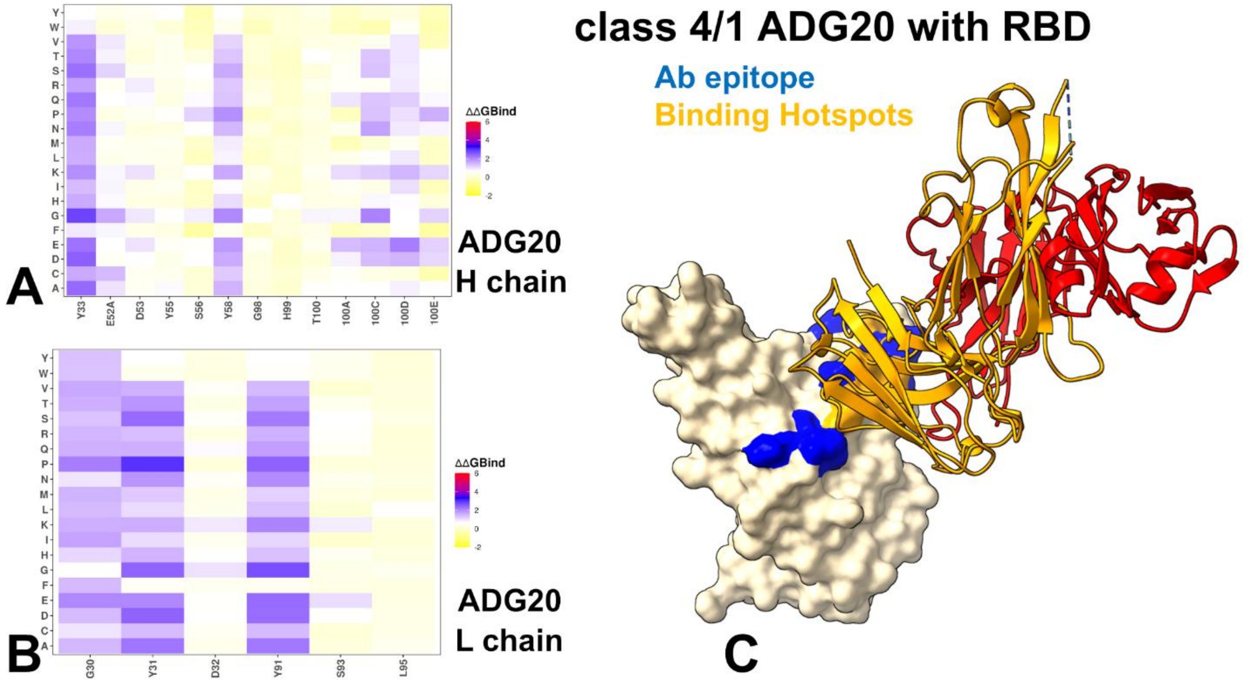


**Figure S7. Ensemble-based dynamic mutational profiling of the RBD intermolecular interfaces in the RBD complex with ADG20 antibody.** The mutational scanning heatmaps are shown for the interfacial heavy chain residues of ADG20 (A) and light chain residues of ADG20 (B). The heatmaps show the computed binding free energy changes for 20 single mutations of the interfacial positions. (C) The structure of ADG20 bound to RBD. The heavy chain of ADG20 is in orange ribbons, and light chain is in red ribbons. The binding epitope is shown in blue surface and the positions of the RBD binding energy hotspots are shown in orange-colored surface. (D). RBD from the complex with ADG20. The binding epitope residues are in blue surface and the binding interfacial RBD hotspots are in orange surface.

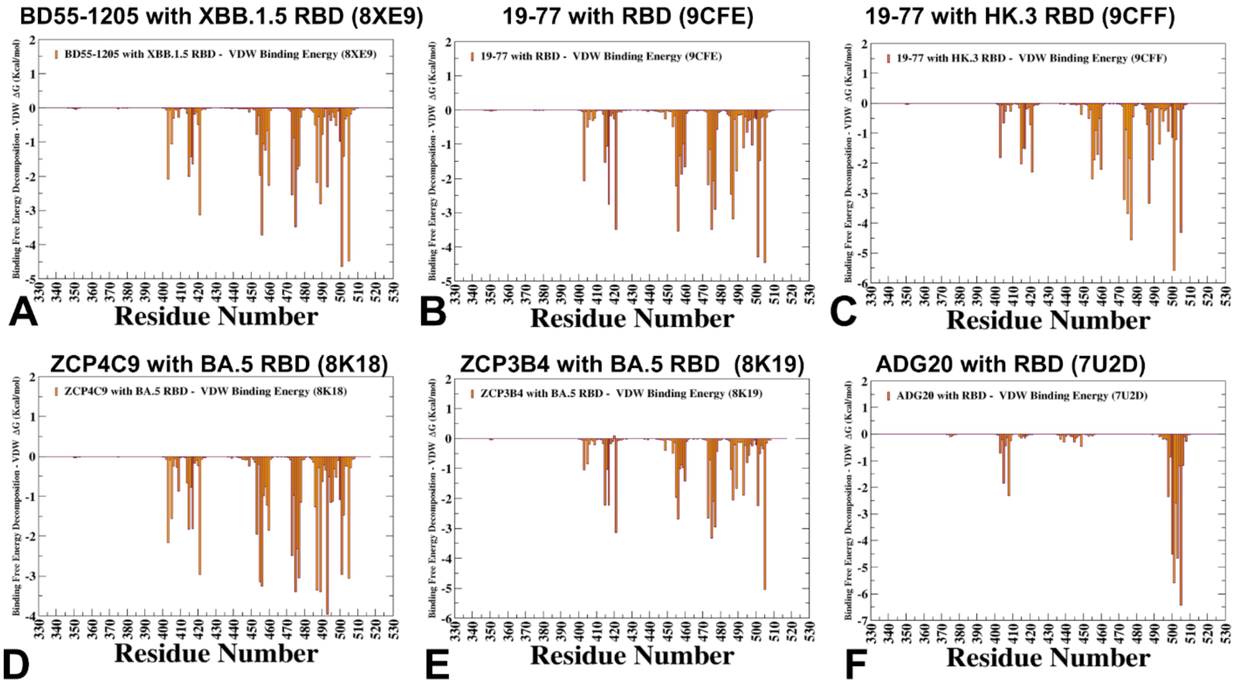


**Figure S8.** The residue-based decomposition of the van der Waals contribution of MM-GBSA energies for the S-RBD complexes with class 1 BD55-1205 with XBB.1.5 RBD (A), 19-77 class 1 antibody with RBD (B), 19-77 class 1 antibody with HK.3 RBD (C), class 1 ZCP4C9 with

BA.5 RBD (D), class 1 ZCP3B4 with BA.5 RBD (E), class 4/1 antibody ADG20 with RBD (F). The binding free energy with MM-GBSA was computed by averaging the results of computations over 10,000 samples from the equilibrium ensembles.

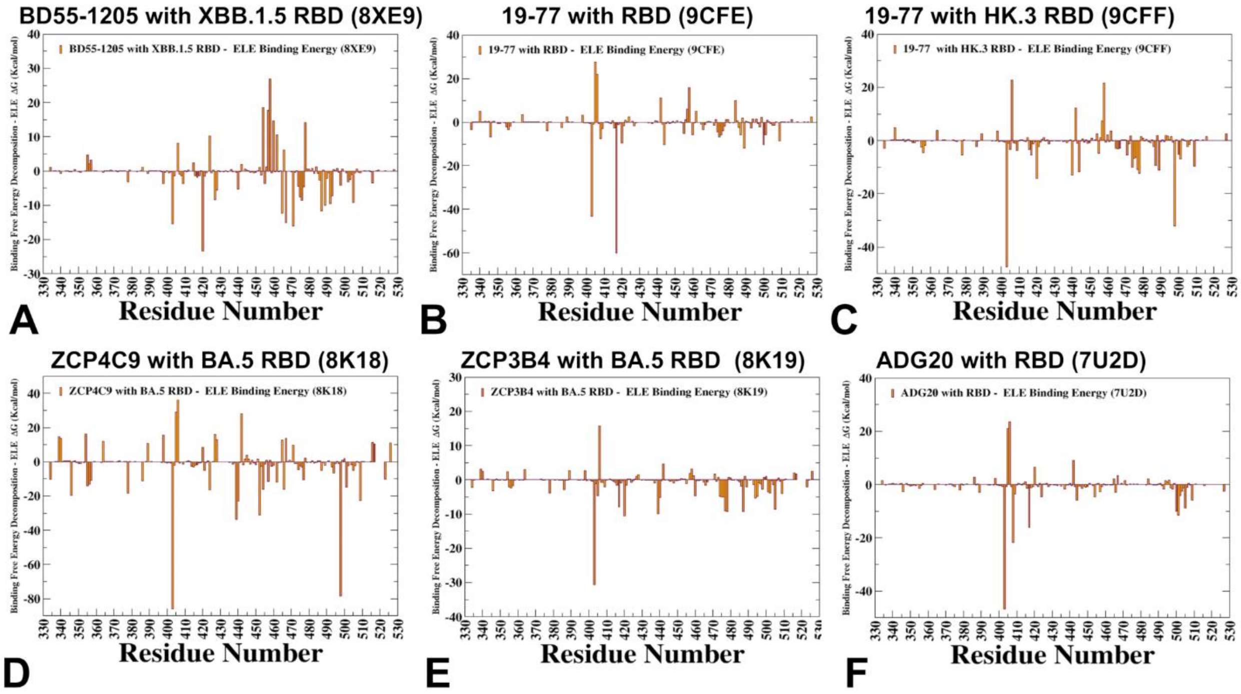


**Figure S9.** The residue-based decomposition of the electrostatic contribution of MM-GBSA energies for the S-RBD complexes with class 1 BD55-1205 with XBB.1.5 RBD (A), 19-77 class 1 antibody with RBD (B), 19-77 class 1 antibody with HK.3 RBD (C), class 1 ZCP4C9 with BA.5 RBD (D), class 1 ZCP3B4 with BA.5 RBD (E), class 4/1 antibody ADG20 with RBD (F). The binding free energy with MM-GBSA was computed by averaging the results of computations over 10,000 samples from the equilibrium ensembles.

**Table S1.** Mutational landscape of the Omicron variants.

| Variant | Mutational landscape |
|---|---|
| XBB.1.5 | T19I, V83A, G142D, Del144, H146Q, Q183E, V213E, G252V, G339H, R346T, L368I, S371F, S373P, S375F, T376A, D405N, R408S, K417N, N440K, V445P, G446S,N460K, S477N, T478K, E484A, **F486P, F490S**, R493Q reversal, Q498R, N501Y, Y505H, D614G, H655Y, N679K, P681H, N764K, D796Y, Q954H, N969K |
| JN.1 | T19I, R21T, S50L, del69-70,V127F, delY144, F157S, R158G, delN211, L213I, L226F, H25N, A264D, I332V, D339H, K356T, R403K, V445H, G446S, N450D, L452W, **L455S,** N460K, N481K, del V483, A484K, F486P, R493Q, E554K, A570V, P612S, I670V, H68R, D939F, P1143L |
| KP.2 | **JN.1 + S:R346T, S:F456L, S:V1104L**<br><br>T19I, R21T, S50L, del69-70,V127F, delY144, F157S, R158G, delN211, L213I, L226F, H25N,A264D, I332V, D339H, **R346T**, K356T, R403K, V445H, G446S, N450D, L452W, **L455S, F456L**, N460K, N481K, del V483, A484K, F486P, R493Q, E554K, A570V, P612S, I670V, H68R, D939F, **V1104L,** P1143L |
| KP.3 | **JN.1 + S:F456L, S:Q493E, S:V1104L**<br><br>T19I, R21T, S50L, del69-70,V127F, delY144, F157S, R158G, delN211, L213I, L226F, H25N,A264D, I332V, D339H, K356T, R403K, V445H, G446, N450D, L452W, **L455S, F456L**, N460K, N481K, del V483, A484K, F486P, **Q493E**, E554K, A570V, P612S, I670V, H68R, D939F, **V1104L,** P1143L |
| KP.1.1 | **JN.1 + S:F456L, S:R346T, S:K1086R, S:V1104L**<br><br>T19I, R21T, S50L, del69-70,V127F, delY144, F157S, R158G, delN211, L213I, L226F, H25N,A264D, I332V, D339H, **R346T**, K356T, R403K, V445H, G446, N450D, L452W, **L455S, F456L**, N460K, N481K, del V483, A484K, F486P, **Q493E**, E554K, A570V, P612S, H68R, D939F, **K1086R**, **V1104L,** P1143L |

| | |
|---|---|
| KP.3.1.1 | **KP.3 + S:S31-**<br><br>T19I, R21T, S31-, S50L, del69-70,V127F, delY144, F157S, R158G, delN211, L213I, L226F, H25N,A264D, I332V, D339H, K356T, R403K, V445H, G446, N450D, L452W, L455S, F456L, N460K, N481K, del V483, A484K, F486P, Q493E, E554K, A570V, P612S, I670V, H68R, D939F, V1104L, P1143L |
| LP.8 | **KP.1.1+ F186L, H445R, Q493E, S31 del**<br><br>T19I, R21T, S31 del, S50L, del69-70,V127F, delY144, F157S, R158G, F186L, delN211, L213I, L226F, H25N,A264D, I332V, D339H, **R346T**, K356T, R403K, H445R, G446, N450D, L452W, **L455S, F456L**, N460K, N481K, del V483, A484K, F486P, **Q493E**, E554K, A570V, P612S, I670V, H68R, D939F, **K1086R**, **V1104L,** P1143L |
| LB.1 | **JN.1+ S:S31-, S:Q183H, S:R346T, S:F456L**<br><br>T19I, R21T, **S31-**, S50L, del69-70,V127F, delY144, F157S, R158G, **Q183H**, delN211, L213I, L226F, H25N,A264D, I332V, D339H, **R346T,** K356T, R403K, V445H, G446S, N450D, L452W, **L455S, F456L**, N460K, N481K, del V483, A484K, F486P, R493Q, E554K, A570V, P612S, I670V, H68R, D939F, P1143L |
| XEC | **JN.1 + S:T22N, S:F59S, S:F456L, S:Q493E, S:V1104L**<br><br>T19I, R21T, **T22N**, S50L, **F59S**, del69-70,V127F, delY144, F157S, R158G, delN211, L213I, L226F, H25N,A264D, I332V, D339H, K356T, R403K, V445H, G446S, N450D, L452W, **L455S,** F456L. N460K, N481K, del V483, A484K, F486P, **Q493E**, E554K, A570V, P612S, I670V, H68R, D939F, **V1104L**, P1143L |
| LB.8.1 | **JN1 + S:S31-, S:F186L, S:R190S, S:R346T, S:V445R, S:F456L, S:Q493E, S:K1086R, S:V1104L**<br><br>T19I, R21T, S31-, S50L, del69-70,V127F, delY144, F157S, R158G, F186L, R190S, delN211, L213I, L226F, H25N, A264D, I332V, D339H, R346T, K356T, R403K, V445R, G446S, N450D, L452W, **L455S,** F456L, N460K, N481K, del V483, A484K, F486P, Q493E, E554K, A570V, P612S, I670V, H68R, D939F, K1086R, V1140L, P1143L |
| NB.1.8.1 | **JN1 + S:T22N, S:F59S, S:G184S,S:A435S, S:F456L, S:T478I, S:Q493E** |
| XFG | **JN1 + S:T22N, S:S31P, S:K182R, S:R190S, S:R346T, S:K444R, S:V445R, S:F456L, S:N487D, S:Q493E, S:T572I** |

**Table S2.** The list of the intermolecular contacts in the structure of the Class 1 BD55-1205 antibody complex with XBB.1.5 RBD (pdb id 8XE9). The interfacial contacts in the structure are defined by counting the number of interatomic contacts within a 5.5 Å distance threshold between atoms of the interacting proteins.

| RBD Residue | RBD Residue Number | RBD chain | Ab Residue | Ab Residue Number | Ab chain |
|---|---|---|---|---|---|
| ARG | 403 | C | ASN | 30 | B |
| ARG | 403 | C | GLY | 92 | B |
| ASN | 405 | C | ASP | 93 | B |
| THR | 415 | C | SER | 56 | A |
| THR | 415 | C | THR | 57 | A |
| THR | 415 | C | PHE | 58 | A |
| GLY | 416 | C | TYR | 52 | A |
| GLY | 416 | C | SER | 56 | A |
| GLY | 416 | C | PHE | 58 | A |
| ASN | 417 | C | TYR | 33 | A |
| ASN | 417 | C | TYR | 52 | A |
| ASN | 417 | C | TRP | 94 | B |
| ASN | 417 | C | PRO | 95 | B |
| ASP | 420 | C | TYR | 52 | A |
| ASP | 420 | C | SER | 56 | A |
| TYR | 421 | C | TYR | 33 | A |
| TYR | 421 | C | TYR | 52 | A |
| TYR | 421 | C | PRO | 53 | A |
| TYR | 421 | C | GLY | 54 | A |
| TYR | 421 | C | GLY | 55 | A |
| TYR | 453 | C | ILE | 101 | A |
| LEU | 455 | C | TYR | 33 | A |
| LEU | 455 | C | PRO | 53 | A |
| LEU | 455 | C | TRP | 94 | B |

| | | | | | |
|---|---|---|---|---|---|
| LEU | 455 | C | LEU | 99 | A |
| LEU | 455 | C | ILE | 101 | A |
| LEU | 455 | C | ARG | 102 | A |
| PHE | 456 | C | ARG | 31 | A |
| PHE | 456 | C | ASN | 32 | A |
| PHE | 456 | C | TYR | 33 | A |
| PHE | 456 | C | PRO | 53 | A |
| PHE | 456 | C | LEU | 99 | A |
| ARG | 457 | C | PRO | 53 | A |
| ARG | 457 | C | GLY | 54 | A |
| LYS | 458 | C | SER | 30 | A |
| LYS | 458 | C | ARG | 31 | A |
| LYS | 458 | C | PRO | 53 | A |
| LYS | 458 | C | GLY | 54 | A |
| SER | 459 | C | PRO | 53 | A |
| SER | 459 | C | GLY | 54 | A |
| LYS | 460 | C | GLY | 54 | A |
| LYS | 460 | C | GLY | 55 | A |
| LYS | 460 | C | SER | 56 | A |
| TYR | 473 | C | SER | 30 | A |
| TYR | 473 | C | ARG | 31 | A |
| TYR | 473 | C | ASN | 32 | A |
| TYR | 473 | C | PRO | 53 | A |
| GLN | 474 | C | ARG | 31 | A |
| ALA | 475 | C | PHE | 27 | A |
| ALA | 475 | C | THR | 28 | A |
| ALA | 475 | C | ARG | 31 | A |
| ALA | 475 | C | ASN | 32 | A |
| ALA | 475 | C | ARG | 97 | A |
| GLY | 476 | C | GLY | 26 | A |
| GLY | 476 | C | PHE | 27 | A |
| GLY | 476 | C | THR | 28 | A |
| GLY | 476 | C | ARG | 31 | A |
| GLY | 476 | C | ASN | 32 | A |
| ASN | 477 | C | GLY | 26 | A |
| ASN | 477 | C | PHE | 27 | A |
| ASN | 477 | C | THR | 28 | A |
| PRO | 486 | C | GLU | 104 | A |
| ASN | 487 | C | VAL | 2 | A |
| ASN | 487 | C | GLY | 26 | A |
| ASN | 487 | C | PHE | 27 | A |
| ASN | 487 | C | ARG | 97 | A |
| ASN | 487 | C | GLU | 104 | A |

| | | | | | |
|---|---|---|---|---|---|
| TYR | 489 | C | ASN | 32 | A |
| TYR | 489 | C | ARG | 97 | A |
| TYR | 489 | C | LEU | 99 | A |
| TYR | 489 | C | ARG | 102 | A |
| TYR | 489 | C | GLU | 104 | A |
| SER | 490 | C | ARG | 102 | A |
| PRO | 491 | C | ARG | 102 | A |
| LEU | 492 | C | ARG | 102 | A |
| GLN | 493 | C | ILE | 101 | A |
| GLN | 493 | C | ARG | 102 | A |
| ARG | 498 | C | SER | 31 | B |
| ARG | 498 | C | SER | 67 | B |
| THR | 500 | C | SER | 28 | B |
| THR | 500 | C | PHE | 29 | B |
| THR | 500 | C | GLY | 68 | B |
| TYR | 501 | C | SER | 28 | B |
| TYR | 501 | C | PHE | 29 | B |
| TYR | 501 | C | ASN | 30 | B |
| TYR | 501 | C | SER | 31 | B |
| GLY | 502 | C | SER | 28 | B |
| GLY | 502 | C | PHE | 29 | B |
| GLY | 502 | C | ASN | 30 | B |
| VAL | 503 | C | SER | 28 | B |
| HIS | 505 | C | SER | 28 | B |
| HIS | 505 | C | PHE | 29 | B |
| HIS | 505 | C | ASN | 30 | B |
| HIS | 505 | C | GLY | 92 | B |
| HIS | 505 | C | ASP | 93 | B |

**Table S3.** The list of the intermolecular contacts in the structure of the Class 1 19-77 antibody complex with RBD (pdb id 9CFE). The interfacial contacts in the structure are defined by counting the number of interatomic contacts within a 5.5 Å distance threshold between atoms of the interacting proteins.

| RBD Residue | RBD Residue Number | RBD chain | Ab Residue | Ab Residue Number | Ab chain |
|---|---|---|---|---|---|
| ARG | 403 | C | PHE | 32 | B |
| ARG | 403 | C | SER | 92 | B |
| ARG | 403 | C | ASP | 93 | B |
| ASP | 405 | C | SER | 92 | B |
| ARG | 408 | C | PHE | 58 | A |
| THR | 415 | C | SER | 56 | A |
| THR | 415 | C | PHE | 58 | A |
| GLY | 416 | C | TYR | 52 | A |
| GLY | 416 | C | SER | 56 | A |
| GLY | 416 | C | PHE | 58 | A |
| LYS | 417 | C | TYR | 33 | A |
| LYS | 417 | C | TYR | 52 | A |
| LYS | 417 | C | ASP | 93 | B |
| LYS | 417 | C | TRP | 94 | B |
| LYS | 417 | C | PRO | 95 | B |
| ASP | 420 | C | TYR | 52 | A |
| ASP | 420 | C | SER | 56 | A |
| ASP | 420 | C | PHE | 58 | A |
| TYR | 421 | C | TYR | 33 | A |
| TYR | 421 | C | TYR | 52 | A |
| TYR | 421 | C | PRO | 53 | A |
| TYR | 421 | C | GLY | 54 | A |
| TYR | 421 | C | GLY | 55 | A |
| TYR | 421 | C | SER | 56 | A |

| | | | | | |
|---|---|---|---|---|---|
| LEU | 455 | C | TYR | 33 | A |
| LEU | 455 | C | PRO | 53 | A |
| LEU | 455 | C | TRP | 94 | B |
| LEU | 455 | C | PRO | 100 | A |
| PHE | 456 | C | ARG | 31 | A |
| PHE | 456 | C | ASN | 32 | A |
| PHE | 456 | C | TYR | 33 | A |
| PHE | 456 | C | PRO | 53 | A |
| PHE | 456 | C | TRP | 94 | B |
| PHE | 456 | C | ASP | 98 | A |
| PHE | 456 | C | ALA | 99 | A |
| PHE | 456 | C | PRO | 100 | A |
| ARG | 457 | C | ARG | 31 | A |
| ARG | 457 | C | PRO | 53 | A |
| ARG | 457 | C | GLY | 54 | A |
| LYS | 458 | C | SER | 30 | A |
| LYS | 458 | C | ARG | 31 | A |
| LYS | 458 | C | PRO | 53 | A |
| LYS | 458 | C | GLY | 54 | A |
| LYS | 458 | C | ARG | 71 | A |
| SER | 459 | C | PRO | 53 | A |
| SER | 459 | C | GLY | 54 | A |
| ASN | 460 | C | PRO | 53 | A |
| ASN | 460 | C | GLY | 54 | A |
| ASN | 460 | C | GLY | 55 | A |
| ASN | 460 | C | SER | 56 | A |
| TYR | 473 | C | SER | 30 | A |
| TYR | 473 | C | ARG | 31 | A |
| TYR | 473 | C | ASN | 32 | A |
| TYR | 473 | C | PRO | 53 | A |
| GLN | 474 | C | ILE | 28 | A |
| GLN | 474 | C | ARG | 31 | A |
| ALA | 475 | C | GLU | 26 | A |
| ALA | 475 | C | LEU | 27 | A |
| ALA | 475 | C | ILE | 28 | A |
| ALA | 475 | C | ARG | 31 | A |
| ALA | 475 | C | ASN | 32 | A |
| ALA | 475 | C | ARG | 97 | A |
| GLY | 476 | C | GLU | 26 | A |
| GLY | 476 | C | LEU | 27 | A |
| GLY | 476 | C | ILE | 28 | A |
| GLY | 476 | C | ASN | 32 | A |
| SER | 477 | C | SER | 25 | A |

| | | | | | |
|---|---|---|---|---|---|
| SER | 477 | C | GLU | 26 | A |
| SER | 477 | C | LEU | 27 | A |
| SER | 477 | C | ILE | 28 | A |
| THR | 478 | C | GLU | 26 | A |
| GLY | 485 | C | GLU | 102 | A |
| PHE | 486 | C | GLU | 1 | A |
| PHE | 486 | C | VAL | 2 | A |
| PHE | 486 | C | GLU | 26 | A |
| PHE | 486 | C | GLU | 102 | A |
| ASN | 487 | C | VAL | 2 | A |
| ASN | 487 | C | GLU | 26 | A |
| ASN | 487 | C | LEU | 27 | A |
| ASN | 487 | C | ARG | 97 | A |
| ASN | 487 | C | GLU | 102 | A |
| ASN | 487 | C | ASP | 104 | A |
| TYR | 489 | C | ASN | 32 | A |
| TYR | 489 | C | ARG | 97 | A |
| TYR | 489 | C | ALA | 99 | A |
| TYR | 489 | C | PRO | 100 | A |
| TYR | 489 | C | SER | 101 | A |
| TYR | 489 | C | GLU | 102 | A |
| GLN | 493 | C | ASP | 50 | B |
| GLN | 493 | C | PRO | 100 | A |
| TYR | 495 | C | HIS | 31 | B |
| TYR | 495 | C | PHE | 32 | B |
| GLY | 496 | C | HIS | 31 | B |
| GLN | 498 | C | HIS | 31 | B |
| PRO | 499 | C | GLU | 68 | B |
| THR | 500 | C | ASN | 28 | B |
| THR | 500 | C | GLU | 68 | B |
| ASN | 501 | C | ASN | 28 | B |
| ASN | 501 | C | ILE | 29 | B |
| ASN | 501 | C | GLY | 30 | B |
| ASN | 501 | C | HIS | 31 | B |
| ASN | 501 | C | SER | 67 | B |
| ASN | 501 | C | GLU | 68 | B |
| GLY | 502 | C | ASN | 28 | B |
| GLY | 502 | C | GLY | 30 | B |
| GLY | 502 | C | GLU | 68 | B |
| TYR | 505 | C | ILE | 2 | B |
| TYR | 505 | C | GLN | 27 | B |
| TYR | 505 | C | ASN | 28 | B |
| TYR | 505 | C | ILE | 29 | B |

| | | | | | |
|---|---|---|---|---|---|
| TYR | 505 | C | GLY | 30 | B |
| TYR | 505 | C | PHE | 32 | B |
| TYR | 505 | C | GLU | 90 | B |
| TYR | 505 | C | SER | 92 | B |

**Table S4.** The list of the intermolecular contacts in the structure of the Class 1 ZCP4C9 antibody complex with BA.5 RBD (pdb id 8K18). The interfacial contacts in the structure are defined by counting the number of interatomic contacts within a 5.5 Å distance threshold between atoms of the interacting proteins.

| RBD Residue | RBD Residue Number | RBD chain | Ab Residue | Ab Residue Number | Ab chain |
|---|---|---|---|---|---|
| LYS | 403 | E | ASN | 30 | D |
| LYS | 403 | E | TRP | 102 | C |
| ASP | 405 | E | GLY | 92 | D |
| ASP | 405 | E | HIS | 93 | D |
| SER | 408 | E | GLN | 94 | D |
| GLN | 409 | E | GLN | 94 | D |
| GLN | 414 | E | GLN | 94 | D |
| THR | 415 | E | SER | 56 | C |
| THR | 415 | E | THR | 57 | C |
| THR | 415 | E | PHE | 58 | C |
| GLY | 416 | E | SER | 56 | C |
| GLY | 416 | E | PHE | 58 | C |
| GLY | 416 | E | GLN | 94 | D |
| ASN | 417 | E | TYR | 33 | C |
| ASN | 417 | E | TYR | 52 | C |
| ASN | 417 | E | PHE | 58 | C |
| ASP | 420 | E | GLY | 55 | C |
| ASP | 420 | E | SER | 56 | C |
| TYR | 421 | E | TYR | 33 | C |
| TYR | 421 | E | TYR | 52 | C |
| TYR | 421 | E | PRO | 53 | C |
| TYR | 421 | E | GLY | 54 | C |
| TYR | 421 | E | GLY | 55 | C |
| TYR | 421 | E | SER | 56 | C |
| TYR | 453 | E | TRP | 102 | C |

| | | | | | |
|---|---|---|---|---|---|
| TYR | 453 | E | TYR | 104 | C |
| LEU | 455 | E | TYR | 33 | C |
| LEU | 455 | E | PRO | 53 | C |
| LEU | 455 | E | LEU | 99 | C |
| LEU | 455 | E | GLU | 101 | C |
| LEU | 455 | E | TYR | 104 | C |
| PHE | 456 | E | ARG | 31 | C |
| PHE | 456 | E | TYR | 33 | C |
| PHE | 456 | E | PRO | 53 | C |
| PHE | 456 | E | LEU | 99 | C |
| PHE | 456 | E | LEU | 100 | C |
| ARG | 457 | E | PRO | 53 | C |
| ARG | 457 | E | GLY | 54 | C |
| LYS | 458 | E | ARG | 31 | C |
| LYS | 458 | E | PRO | 53 | C |
| LYS | 458 | E | GLY | 54 | C |
| SER | 459 | E | PRO | 53 | C |
| SER | 459 | E | GLY | 54 | C |
| ASN | 460 | E | GLY | 54 | C |
| ASN | 460 | E | GLY | 55 | C |
| ASN | 460 | E | SER | 56 | C |
| TYR | 473 | E | SER | 30 | C |
| TYR | 473 | E | ARG | 31 | C |
| TYR | 473 | E | ASN | 32 | C |
| TYR | 473 | E | PRO | 53 | C |
| TYR | 473 | E | GLY | 54 | C |
| GLN | 474 | E | ARG | 31 | C |
| ALA | 475 | E | ILE | 28 | C |
| ALA | 475 | E | VAL | 29 | C |
| ALA | 475 | E | SER | 30 | C |
| ALA | 475 | E | ARG | 31 | C |
| ALA | 475 | E | ASN | 32 | C |
| GLY | 476 | E | PHE | 27 | C |
| GLY | 476 | E | ILE | 28 | C |
| GLY | 476 | E | SER | 30 | C |
| GLY | 476 | E | ARG | 31 | C |
| GLY | 476 | E | ASN | 32 | C |
| ASN | 477 | E | ILE | 28 | C |
| ASN | 477 | E | SER | 30 | C |
| ASN | 477 | E | ARG | 31 | C |
| ASN | 477 | E | ASN | 76 | C |
| LYS | 478 | E | ILE | 28 | C |
| VAL | 486 | E | GLY | 26 | C |

| | | | | | |
|---|---|---|---|---|---|
| ASN | 487 | E | GLY | 26 | C |
| ASN | 487 | E | PHE | 27 | C |
| ASN | 487 | E | ILE | 28 | C |
| ASN | 487 | E | ASN | 32 | C |
| TYR | 489 | E | PHE | 27 | C |
| TYR | 489 | E | ASN | 32 | C |
| TYR | 489 | E | ARG | 97 | C |
| TYR | 489 | E | GLY | 98 | C |
| TYR | 489 | E | LEU | 99 | C |
| TYR | 489 | E | LEU | 100 | C |
| PHE | 490 | E | LEU | 100 | C |
| GLN | 493 | E | LEU | 99 | C |
| GLN | 493 | E | LEU | 100 | C |
| GLN | 493 | E | GLU | 101 | C |
| GLN | 493 | E | TRP | 102 | C |
| GLN | 493 | E | TYR | 104 | C |
| SER | 494 | E | TRP | 102 | C |
| TYR | 495 | E | TRP | 102 | C |
| SER | 496 | E | GLU | 31 | D |
| SER | 496 | E | TRP | 102 | C |
| ARG | 498 | E | GLU | 31 | D |
| ARG | 498 | E | PHE | 52 | D |
| THR | 500 | E | ASP | 28 | D |
| THR | 500 | E | GLU | 31 | D |
| THR | 500 | E | SER | 67 | D |
| THR | 500 | E | GLY | 68 | D |
| TYR | 501 | E | ASP | 28 | D |
| TYR | 501 | E | ASN | 30 | D |
| TYR | 501 | E | GLU | 31 | D |
| TYR | 501 | E | GLY | 68 | D |
| GLY | 502 | E | ASP | 28 | D |
| GLY | 502 | E | ASN | 30 | D |
| GLY | 502 | E | GLY | 68 | D |
| VAL | 503 | E | ASP | 28 | D |
| GLY | 504 | E | ASP | 28 | D |
| HIS | 505 | E | ASP | 28 | D |
| HIS | 505 | E | VAL | 29 | D |
| HIS | 505 | E | ASN | 30 | D |
| HIS | 505 | E | ASP | 32 | D |
| HIS | 505 | E | GLY | 92 | D |
| HIS | 505 | E | HIS | 93 | D |
| GLN | 506 | E | ASN | 30 | D |

**Table S5.** The list of the intermolecular contacts in the structure of the Class 1 ZCP3B4 antibody complex with BA.5 RBD (pdb id 8K19). The interfacial contacts in the structure are defined by counting the number of interatomic contacts within a 5.5 Å distance threshold between atoms of the interacting proteins.

| RBD Residue | RBD Residue Number | RBD chain | Ab Residue | Ab Residue Number | Ab chain |
|---|---|---|---|---|---|
| LYS | 403 | E | ASP | 92 | B |
| LYS | 403 | E | HIS | 93 | B |
| ASN | 405 | E | HIS | 93 | B |
| ASP | 406 | E | HIS | 93 | B |
| THR | 415 | E | TYR | 52 | A |
| THR | 415 | E | THR | 56 | A |
| THR | 415 | E | THR | 57 | A |
| THR | 415 | E | TYR | 58 | A |
| GLY | 416 | E | TYR | 52 | A |
| GLY | 416 | E | THR | 56 | A |
| GLY | 416 | E | TYR | 58 | A |
| ASN | 417 | E | TYR | 33 | A |
| ASN | 417 | E | TYR | 52 | A |
| ASN | 417 | E | TYR | 58 | A |
| ASP | 420 | E | TYR | 52 | A |
| ASP | 420 | E | THR | 56 | A |
| ASP | 420 | E | TYR | 58 | A |
| TYR | 421 | E | TYR | 33 | A |
| TYR | 421 | E | TYR | 52 | A |
| TYR | 421 | E | PRO | 53 | A |
| TYR | 421 | E | GLY | 54 | A |
| TYR | 421 | E | GLY | 55 | A |
| TYR | 421 | E | MET | 100 | A |
| TYR | 449 | E | ARG | 103 | A |
| TYR | 453 | E | TYR | 32 | B |
| TYR | 453 | E | GLY | 101 | A |

| | | | | | |
|---|---|---|---|---|---|
| TYR | 453 | E | ARG | 103 | A |
| LEU | 455 | E | MET | 100 | A |
| LEU | 455 | E | GLY | 101 | A |
| LEU | 455 | E | GLY | 102 | A |
| PHE | 456 | E | MET | 100 | A |
| PHE | 456 | E | GLY | 101 | A |
| ARG | 457 | E | PRO | 53 | A |
| ARG | 457 | E | GLY | 54 | A |
| ARG | 457 | E | MET | 100 | A |
| LYS | 458 | E | SER | 30 | A |
| LYS | 458 | E | SER | 31 | A |
| LYS | 458 | E | PRO | 53 | A |
| LYS | 458 | E | GLY | 54 | A |
| SER | 459 | E | SER | 30 | A |
| SER | 459 | E | PRO | 53 | A |
| SER | 459 | E | GLY | 54 | A |
| ASN | 460 | E | PRO | 53 | A |
| ASN | 460 | E | GLY | 54 | A |
| ASN | 460 | E | GLY | 55 | A |
| ASN | 460 | E | THR | 56 | A |
| TYR | 473 | E | SER | 30 | A |
| TYR | 473 | E | SER | 31 | A |
| TYR | 473 | E | ASN | 32 | A |
| TYR | 473 | E | PRO | 53 | A |
| TYR | 473 | E | MET | 100 | A |
| GLN | 474 | E | ILE | 28 | A |
| GLN | 474 | E | SER | 31 | A |
| ALA | 475 | E | GLU | 26 | A |
| ALA | 475 | E | ILE | 27 | A |
| ALA | 475 | E | ILE | 28 | A |
| ALA | 475 | E | VAL | 29 | A |
| ALA | 475 | E | SER | 31 | A |
| ALA | 475 | E | ASN | 32 | A |
| ALA | 475 | E | ARG | 97 | A |
| GLY | 476 | E | GLU | 26 | A |
| GLY | 476 | E | ILE | 27 | A |
| GLY | 476 | E | ILE | 28 | A |
| GLY | 476 | E | SER | 31 | A |
| GLY | 476 | E | ASN | 32 | A |
| ASN | 477 | E | GLU | 26 | A |
| ASN | 477 | E | ILE | 27 | A |
| ASN | 477 | E | ILE | 28 | A |
| ASN | 477 | E | ASN | 76 | A |

| | | | | | |
|---|---|---|---|---|---|
| LYS | 478 | E | GLU | 26 | A |
| VAL | 486 | E | VAL | 2 | A |
| VAL | 486 | E | GLU | 26 | A |
| ASN | 487 | E | GLU | 26 | A |
| ASN | 487 | E | ILE | 27 | A |
| TYR | 489 | E | ARG | 97 | A |
| TYR | 489 | E | ILE | 99 | A |
| TYR | 489 | E | HIS | 104 | A |
| TYR | 489 | E | ASP | 107 | A |
| GLN | 493 | E | MET | 100 | A |
| GLN | 493 | E | GLY | 101 | A |
| GLN | 493 | E | GLY | 102 | A |
| GLN | 493 | E | ARG | 103 | A |
| GLN | 493 | E | HIS | 104 | A |
| SER | 494 | E | ARG | 103 | A |
| TYR | 495 | E | ARG | 103 | A |
| SER | 496 | E | ARG | 103 | A |
| TYR | 501 | E | ASP | 28 | B |
| TYR | 501 | E | ASN | 30 | B |
| GLY | 502 | E | ASP | 28 | B |
| VAL | 503 | E | ASP | 28 | B |
| GLY | 504 | E | ASP | 28 | B |
| GLY | 504 | E | HIS | 93 | B |
| HIS | 505 | E | ASP | 28 | B |
| HIS | 505 | E | ILE | 29 | B |
| HIS | 505 | E | ASN | 30 | B |
| HIS | 505 | E | TYR | 32 | B |
| HIS | 505 | E | ASP | 92 | B |
| HIS | 505 | E | HIS | 93 | B |

**Table S6.** The list of the intermolecular contacts in the structure of the Class 4/1 ADG20 antibody complex with BA.5 RBD (pdb id 7U2D). The interfacial contacts in the structure are defined by counting the number of interatomic contacts within a 5.5 Å distance threshold between atoms of the interacting proteins.

| RBD Residue | RBD Residue Number | RBD chain | Ab Residue | Ab Residue Number | Ab chain |
|---|---|---|---|---|---|
| ARG | 403 | A | TYR | 33 | H |
| ASP | 405 | A | TYR | 33 | H |
| ASP | 405 | A | SER | 52 | H |
| ASP | 405 | A | ASP | 53 | H |
| ASP | 405 | A | SER | 56 | H |
| ASP | 405 | A | TYR | 58 | H |
| ARG | 408 | A | ASP | 53 | H |
| ARG | 408 | A | TYR | 55 | H |
| ARG | 408 | A | SER | 56 | H |
| GLN | 409 | A | ASP | 53 | H |
| ASN | 439 | A | TYR | 31 | L |
| TYR | 449 | A | HIS | 99 | H |
| GLY | 496 | A | GLY | 98 | H |
| GLN | 498 | A | GLY | 98 | H |
| GLN | 498 | A | HIS | 99 | H |
| GLN | 498 | A | THR | 100 | H |
| PRO | 499 | A | GLY | 30 | L |
| PRO | 499 | A | TYR | 31 | L |
| THR | 500 | A | GLY | 30 | L |
| THR | 500 | A | TYR | 31 | L |
| THR | 500 | A | ASP | 32 | L |
| THR | 500 | A | TRP | 100 | H |
| ASN | 501 | A | TYR | 31 | L |
| ASN | 501 | A | TYR | 91 | L |

| | | | | | |
|---|---|---|---|---|---|
| ASN | 501 | A | GLY | 98 | H |
| ASN | 501 | A | THR | 100 | H |
| ASN | 501 | A | TRP | 100 | H |
| ASN | 501 | A | GLY | 100 | H |
| GLY | 502 | A | TYR | 31 | L |
| GLY | 502 | A | TYR | 91 | L |
| GLY | 502 | A | TRP | 100 | H |
| GLY | 502 | A | GLY | 100 | H |
| VAL | 503 | A | TYR | 31 | L |
| VAL | 503 | A | TYR | 58 | H |
| VAL | 503 | A | TYR | 91 | L |
| VAL | 503 | A | SER | 93 | L |
| VAL | 503 | A | LEU | 95 | L |
| VAL | 503 | A | GLY | 100 | H |
| GLY | 504 | A | TYR | 33 | H |
| GLY | 504 | A | TYR | 58 | H |
| GLY | 504 | A | TYR | 91 | L |
| TYR | 505 | A | SER | 31 | H |
| TYR | 505 | A | TYR | 32 | H |
| TYR | 505 | A | TYR | 33 | H |
| TYR | 505 | A | PHE | 96 | H |
| TYR | 505 | A | SER | 97 | H |
| TYR | 505 | A | GLY | 98 | H |
| TYR | 505 | A | GLY | 100 | H |
| GLN | 506 | A | TYR | 31 | L |
| GLN | 506 | A | TYR | 91 | L |
| GLN | 506 | A | SER | 93 | L |
